# Perturbation biology links temporal protein changes to drug responses in a melanoma cell line

**DOI:** 10.1101/568758

**Authors:** Elin Nyman, Richard R. Stein, Xiaohong Jing, Weiqing Wang, Benjamin Marks, Ioannis K. Zervantonakis, Anil Korkut, Nicholas P. Gauthier, Chris Sander

## Abstract

Data-driven mathematical modeling of biological systems has enormous potential to understand and predict the interplay between molecular and phenotypic response to perturbation, and provides a rational approach to the nomination of therapies in complex diseases such as cancer. Melanoma is a particularly debilitating disease for which most therapies eventually fail as resistance to chemotherapy and targeted drugs develop. We have previously applied an iterative experimental-computational modeling approach, termed perturbation biology, to predict and test effective drug combinations in melanoma cell lines [1].

In this work, we extend our analysis framework to derive models of temporally-acquired perturbation data that do not require prior knowledge and explicit specification of the targets of individual drugs. Specifically, we characterize the response of the melanoma cell line A2058 to 54 cancer drug combinations at 8 logarithmically spaced time points from 10 minutes to 67 hours. At each time point, 124 antibodies of proteins and phospho-proteins with broad coverage of cancer-related pathways and two phenotypes (cell number and apoptosis) were measured. These data are used to infer interactions in ordinary differential equation-based models that capture temporal aspects of the drug perturbation data. This network representation of drug–protein, protein–protein, and protein–phenotype interactions can be used to identify new logical (not necessarily direct biochemical) interactions. The agreement between the predicted phenotypic response and corresponding data for unseen drug perturbations has a Pearson’s correlation coefficient of 0.79. We further use model predictions to nominate effective combination therapies and perform experimental validation of the highest ranked combinations.

This new data-driven modeling framework is a step forward in perturbation biology as it incorporates the temporal aspect of data. This work therefore opens the door to a new understanding of dynamic drug responses at a molecular level.

## Introduction

Targeted therapies are an important component of precision oncology as these agents – as opposed to standard chemotherapy – aim to counteract specific activating genetic or signaling pathway alterations and often have fewer side effects than conventional cytotoxic chemotherapy. Many targeted therapies have been approved by regulatory agencies for the treatment of various cancers [2]. However, initial response rates are generally not durable and tumors eventually develop resistance.

In melanoma, tumors with the common BRAF V600E/K gain-of-function mutation have been shown to have a remarkable response to drugs that specifically target the mutated protein kinase, such as the RAF inhibitor vemurafenib [3]. However, not all patients with a BRAF V600 mutation respond to targeted therapies, and the patients that do respond often develop resistance after only a few months. There are several known mechanisms of this resistance including primary resistance from loss-of-function mutations in PTEN that increase AKT signaling and reduce apoptosis, and CDK4 mutations and CyclinD1 amplification that promote cell cycle progression [4]. Mechanisms of acquired resistance include reactivation of the MEK/ERK pathway with new mutations in BRAF or NRAS and hyper-activation of receptor tyrosine kinases [4]. There is an urgent need for a more comprehensive understanding of resistant tumor cells in order to identify non-trivial therapeutic opportunities beyond targeting single genes.

We have previously developed a perturbation biology approach that simulates data-driven models to find combination vulnerabilities. These models are derived from measurements of the proteomic and phenotypic response of cancer cells *in vitro* to numerous drug combinations. For example, in a melanoma cell line, we predicted that a combination of a RAF or MEK inhibitor with a bromodomain inhibitor would be effective and synergistic and thereby reduce viability [1]. Such a combination (MEK with bromodomain inhibitor), has been subsequently proposed for a phase I/II clinical trial for small cell lung cancer and solid tumors with mutations in RAS [5]. Our approach has also been useful in identifying a synergistic drug combination (CDK4i with IGF1Ri) in dedifferentiated liposarcoma [6]. In these studies, we measured the protein response at a single time point and were thus not able to directly capture the transient cellular response to the drugs.

We hypothesize that short-term responses to therapeutic intervention, implemented by adjustments in signaling networks, already reflect the shifts in cellular processes ultimately implemented in cellular long-term memory via genetic changes. Typically these long-term adaptations are amino acid changing mutations (‘missense mutations’) or amplification/deletion of DNA fragments (‘copy number changes’). It may therefore be possible to obtain evidence for the dominant resistance mechanisms in response to targeted intervention by observing the short term molecular signalling adaptations. In addition, time-series observations of response provide informative input to parameter inference for dynamic models, such as the one developed here, including the directionality, strength and sign of interactions [7, 8]. In order to study the short-term response to targeted therapies in melanoma, we have now produced time-resolved antibody-based measurements of protein and phospho-protein levels as A2058 cancer cells respond and adapt to drug perturbations. A2058 is resistant to both RAF and MEK inhibitors [9], and therefore represents melanoma patients that would fail one of the standard treatments. The obtained response profiles are used to infer interaction parameters of time-resolved mathematical models, which are selected for sparsity and for the ability to predict left-out data at reasonable accuracy. We simulate the consequences of all possible perturbations to these models when it comes to reduction of cell growth and induction of apoptosis. The results of these simulations are ranked by their predicted effectiveness. The most promising candidates, which are expected to reduce cell growth and increase apoptosis in melanoma, involve targeting the combination of IRS1 and EGFR. We experimentally test the drug combination of NT157 (IRS1 inhibitor) with gefitinib (EGFR inhibitor) in A2058 cells, and find that (1) the drugs alone reduces cell growth substantially at high doses, and (2) the drugs in combination reduces cell growth further than any of the drugs alone.

In contrast to models of cell biological processes based on response data at a single time point (assumed to be steady state), deriving models from time-series data may allow one to better capture the dynamic contributions of molecular signaling processes to complex phenotypes such as cell proliferation and apoptosis. This systems biology paradigm of data-driven predictive dynamic models can be applied to other cancers, especially useful for tumors that are resistant to targeted therapies.

## Results

The goal of this data-driven systems biology study is to identify actionable cellular vulnerabilities and as well as to nominate novel vulnerabilities for future drug development. Conceptually, we generate network models that link protein signaling to phenotype. These models are similar to static textbook pathway diagrams, but have the added benefit of being (1) highly specific for the system under study and (2) based on a well-defined mathematical formalism. In this study we produce dynamic (time-resolved) antibody-based measurements of protein and phospho-protein levels as the melanoma cell line A2058 adapts and responds to drugs (alone and in combination). These molecular and phenotypic data are used to train differential equations models that capture cellular dynamics. Using these models, we predict and optimize the effects of untested molecular perturbations on cell growth and apoptosis (Figure 1).

**Figure 1:**
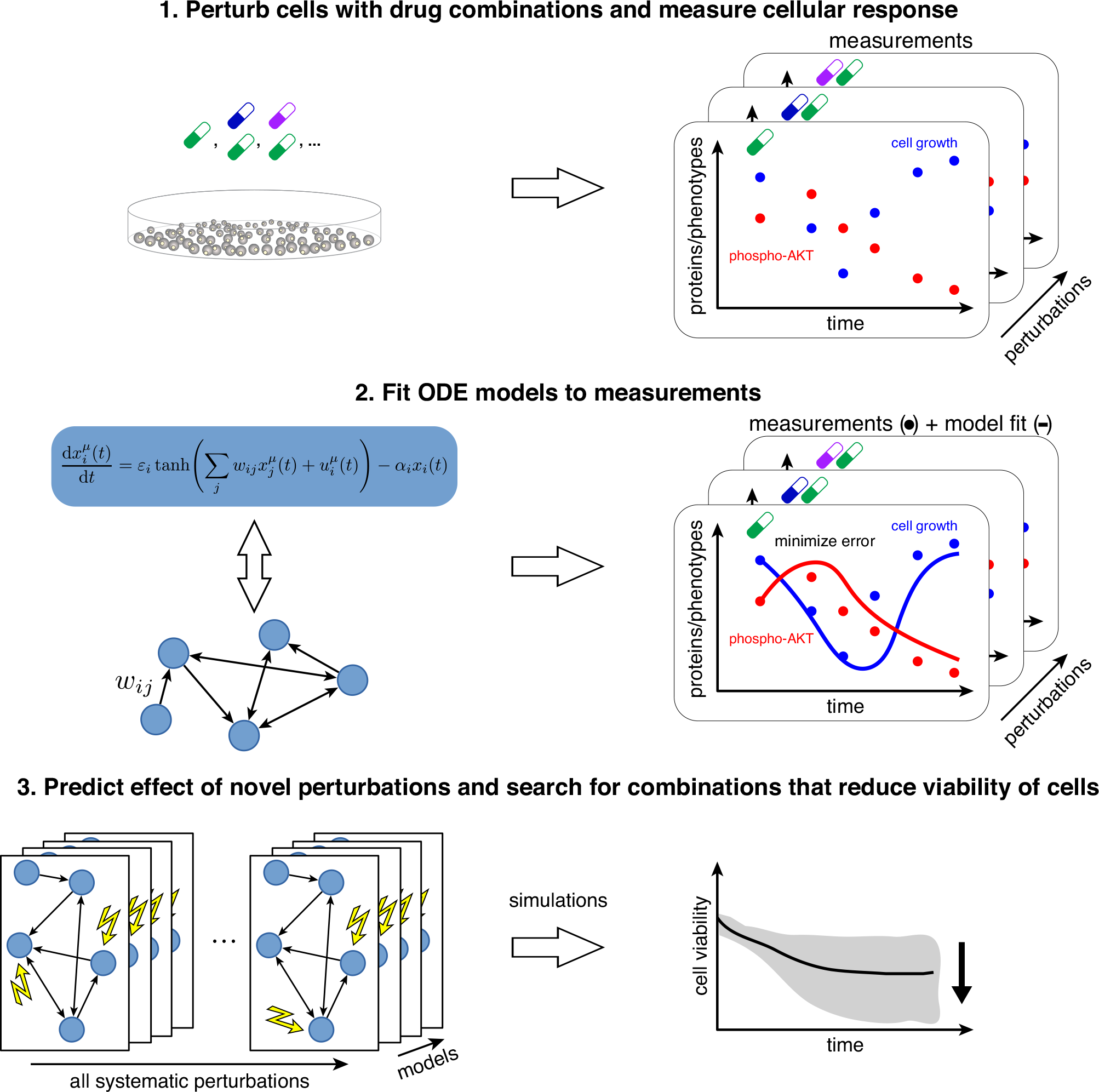
Systematic experimental perturbation of cells leads to predictive network models. We perturb cells with combinations of targeted drugs and measure the time-resolved cellular response (step 1). These measurements are used as input to derive network models of the response to arbitrary combinatorial perturbations (step 2). Using these models, we identify drug combination targets that optimally reduce cell growth and increase apoptosis in a melanoma cell line (step 3).

### Experimental workflow and data

We characterized the temporal response of proteins, phosphorylation sites, cell growth and apoptosis, after application of drugs alone and in combination. A2058 cells were systematically perturbed with as single drugs alone (both low and high doses) or in combination (low doses only), which resulted in 54 total drug conditions (Figure 2).

**Figure 2:**
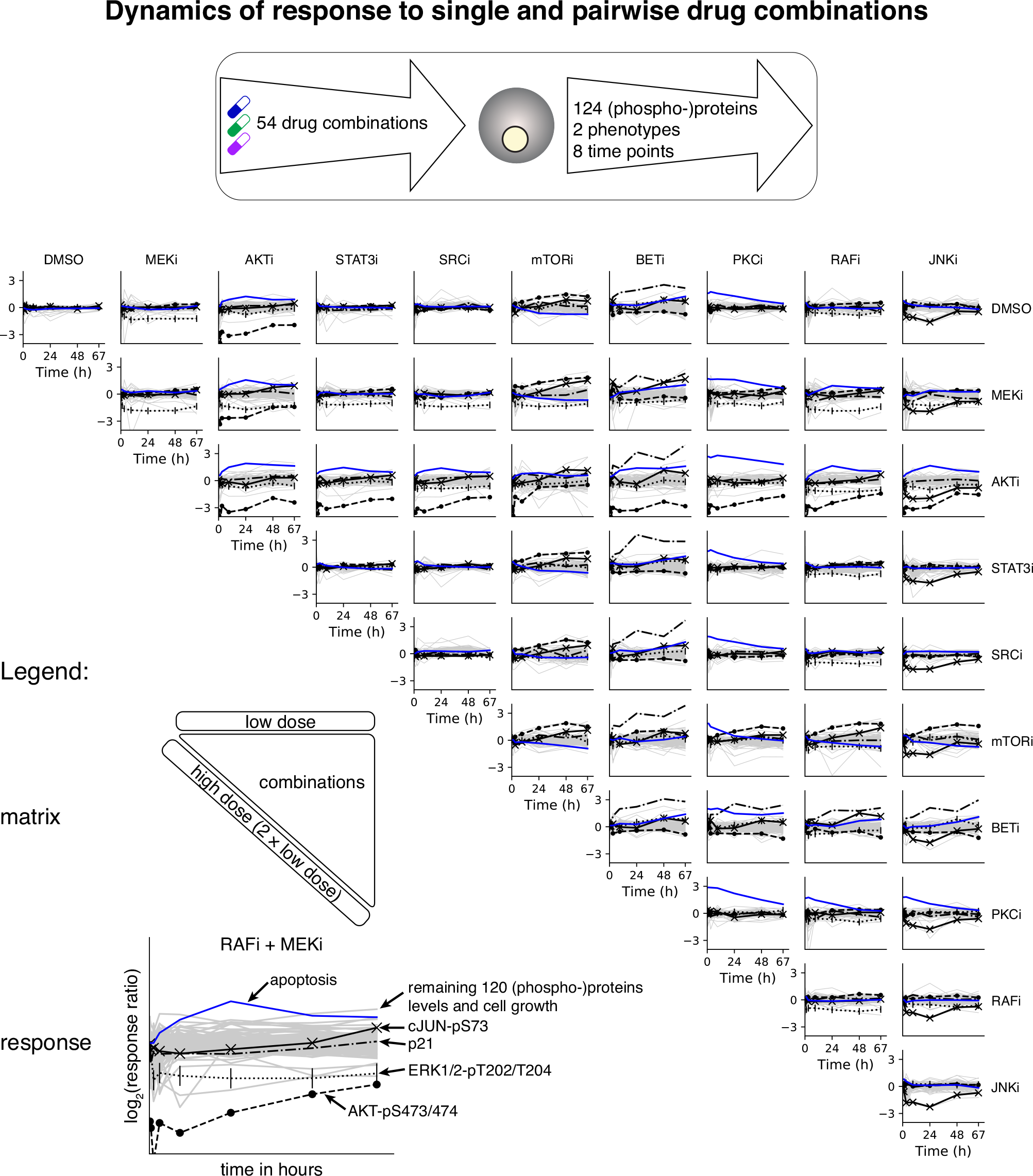
Phenotypic and proteomic response to single and drug combinations. The melanoma cell line A2058 is subjected to 54 drug combinations and the response of 124 (phospho-)proteins and two phenotypes (cell number and apoptosis) is measured at eight time points from 10 minutes to 67 hours. These data are used as input for model inference and prediction. The most informative data involve a subset of the proteins and phospho-proteins (black lines) that undergo the largest changes upon perturbation (AKT-pS473/474, ERK1/2-pT202/T204, cJUN-pS73, and p21) as well as apoptosis (blue line). The remaining 120 proteins and phospho-proteins, as well as cell counts, have a less pronounced response to perturbation. Temporal response (vertical axis) is defined as log_2_ (*x*^perturbed^(*t*)*/x*^unperturbed^(*t*)) where *x*(*t*) are concentrations or counts as in Equations 1 and 2 and DMSO is the unperturbed control.

#### Selection of drug concentrations

For each drug, the concentration was chosen by literature review or experimental dose response measurements by western blot. The *low dose* was selected to reduce activity on a known target by 50%. We also used a *high dose*, which is defined as double the absolute concentration of the *low dose*.

#### Molecular measurements

Cells were collected at 8 time points after drug addition in logarithmic progression (10, 27 and 72 minutes and 3, 9, 24, 48, and 67 hours). The temporal proteomic response of the cells to the different conditions was derived using reverse phase protein array (RPPA) measurements in 124 total and phospho-protein levels [10]. Antibodies were selected to broadly cover signaling pathways with known involvement in cancer (e.g., AKT, ERK, and JAK/STAT pathways).

#### Phenotypic measurements

We used live cell imaging to follow A2058 cells as they responded to drug treatment (Incucyte, Essen BioScience, Ann Arbor, MI, U.S.A.). We acquired GFP, mCherry, and phase images of all conditions every 3 hours for 72 hours following drug addition. Cell number was determined at each time point by counting using image segmentation software (Incucyte), which identified the transgenic H2B-mCherry fluoraphore. Apoptosis was determined by image segmentation and counting of the GFP channel, which measured a Caspase-3/7 fluorescent activity reagent (Essen BioScience). Data were sub-sampled to select time points that closely matched collection of the proteomic data (1, 3, 9, 24, 48, and 67 hours).

#### Agreement of experimental results with literature

As most of the selected drugs have well described signaling effects, i.e., kinase substrates whose activity is directly affected by the drugs (Table 1), we asked whether each drug elicited the expected molecular changes. The drugs that target MEK, AKT, and JNK, resulted in the expected decrease of their known targets ERK1/2-pT202/T204, AKT-pS473/pT308, and CJUN-pS73, respectively (Figure 8). RAF and mTOR inhibition did not result in a particularly strong decrease in direct targets (MEK1/2-pS217/221 and S6K-pT389, respectively), but they did significantly decrease signaling in further downstream nodes ERK1/2-pT202/T204 and S6-pS235/236. For the STAT3 and SRC inhibitors, the proteins’ highest-ranked responders are not known to be downstream targets, but plausibly indirectly affected via multiple steps, signaling feedback or crosstalk.

**Table 1:** Used drugs and their known and herein measured downstream targets.

| Drug | Abbreviation | Downstream targets | Reference |
| --- | --- | --- | --- |
| PD-0325901 | MEKi | ERK1/2-pT202/T204 | [22] |
| AKT1/2i | AKTi | AKT-pS473, AKT-pT308 | [32] |
| Stattic | STAT3i | STAT3-pY705 | [33] |
| CAS 179248-59-0 | SRCi | SRC-pY416, SRC-pY527, CHK2-pT68 | [22] |
| Temsirolimus | mTORi | S6K-pT389, 4EBP1-pS65 | [34, 35] |
| CPI203 | BETi | - |  |
| RO-31-7549 | PKCi | - |  |
| PLX4032 | RAFi | MEK1/2-pS217/221 | [36] |
| JNK-IN-8 | JNKi | CJUN-pS73 | [37] |

When analyzing the phenotypic perturbation readout as measured by live-cell imaging (see Materials and Methods), we found the bromodomain inhibitor CPI203 (BET inhibitor) to be the most potent drug, at the chosen concentrations, in terms of both decrease in cell growth and increase in apoptosis after three days of treatment (Table 1, Figure 7). These data are in agreement with results on the effect of this inhibitor class in the drug-resistant melanoma cell line SkMel-133, which also strongly impaired cell growth in response to the bromodomain inhibitor JQ1 in combination with MEK and ERK inhibitors in our earlier work [1].

### Network model construction and simulation

To analyze the dynamics of molecular and phenotypic response to drug perturbation and to predict the effect of unseen perturbations, we infer and simulate network models from our rich experimental data (Figure 1). We first inferred the parameters for network models on a subset of data (responses to single drugs, training dataset). The optimal level of regularization was then determined on a second subset of data (validation dataset). Finally, the model accuracy was estimated on a third dataset (test dataset). See top left of figure 3. For application purposes, we then derive multiple models on the full dataset to predict the effect of unseen perturbations and rank them by their desired effect on cellular phenotype.

**Figure 3:**
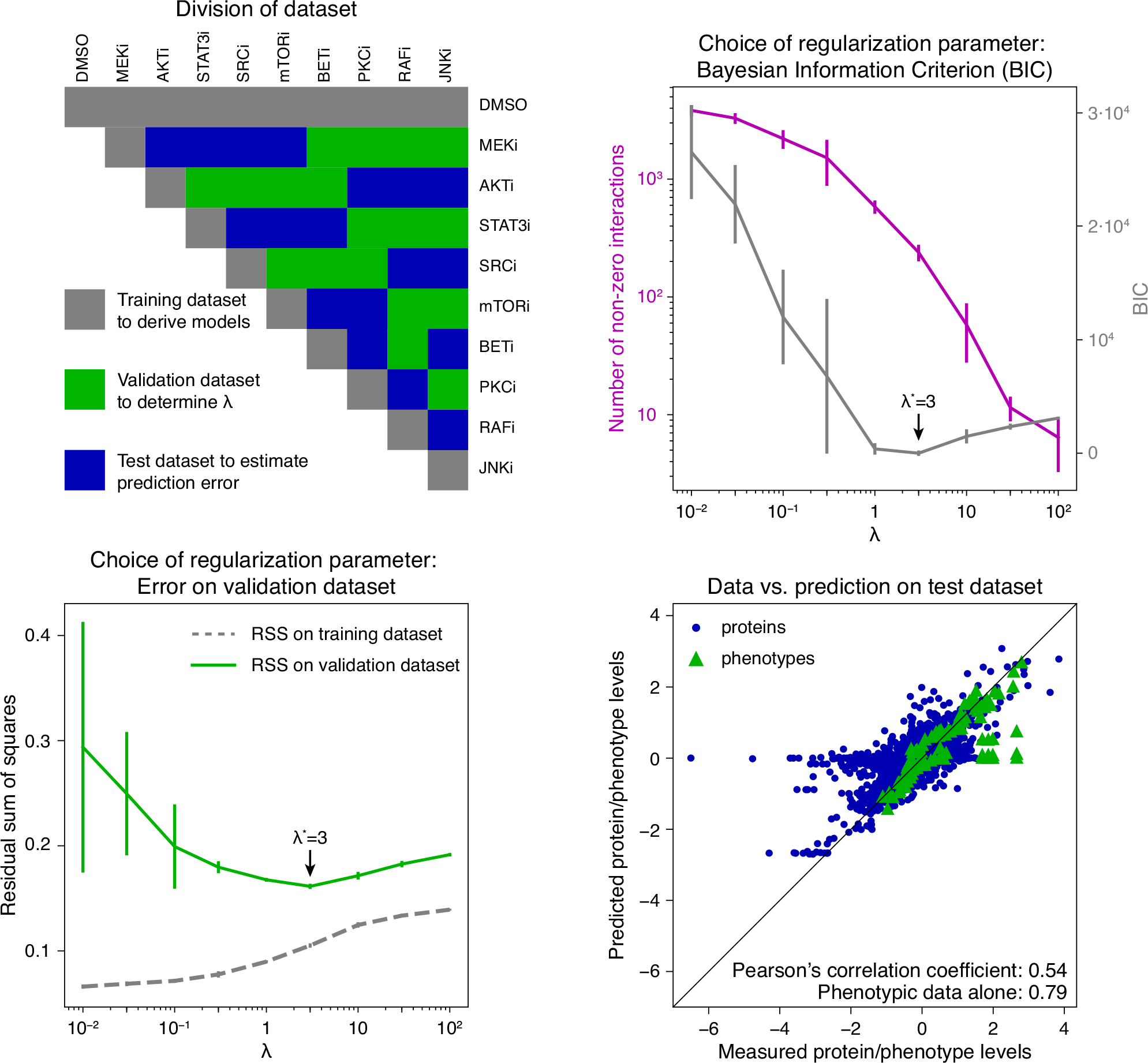
Model selection and error estimation. Top left: The full dataset is divided into subsets: (i) a training dataset (gray) that contains single drug control measurements (DMSO), single drugs in low (first row) and high dose (diagonal, two times low dose), (ii) a validation dataset (green) is used to estimate the optimal regularization parameter *λ*\*, and (iii) a test dataset (blue) is used to estimate model performance. Top right: calculated values for the Bayesian Information Criterion (BIC, gray) and number of non-zero interactions (magenta) as a function of the regularization parameter *λ*. Bottom left: The residual sum of squares on the validation dataset is used to identify the optimal regularization parameter *λ*\*. The best predictive model is obtained for *λ*\* = 3 according to lowest BIC and minimal error on the validation dataset. Error bars indicate the standard deviation from 10 independent runs. Bottom right: Agreement of measured and predicted protein and phospho-protein (dots) and phenotype levels (triangles) on the test dataset. The Pearson correlation coefficient for the combined set of molecular and phenotype nodes is 0.54, and 0.79 for phenotypes alone.

#### Network models

To model the cellular processes, we use the previously proposed nonlinear multiple input–multiple output model [1, 11, 12]. This model has the ability to capture a wide range of biological and kinetic effects, and has been applied to diverse biological problems including to predict the effect of novel drug combinations [1, 6].

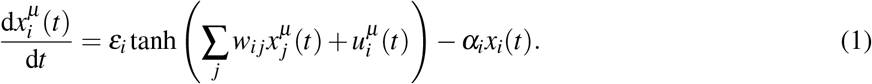

The time-dependent variable 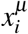 describes the dynamics of node *i* (in our context a protein level, measured phenotype or effect of a drug) and 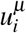 the impact of a perturbation on node *i* (e.g., the application of a single drugs or drug combinations) in experimental condition *μ*. Here, *w*_*i j*_ denotes the interaction between the nodes *i* and *j* (more specifically, molecules, phenotypes, drugs or processes). Intuitively, *w*_*i j*_ > 0 corresponds to activation of node i by node i, and *w*_*i j*_ < 0 to inhibition. The prefactors *α*_*i*_, *ε*_*i*_ > 0 quantify the speed at which the system returns to its (trivial) steady state *x*_*i*_ = 0 and determine the dynamic range of node *i*. The sigmoidal transfer function tanh is used to cap the reaction rates and to account for saturation and noise. To obtain data-derived network models, we used Equation 1 and constrained its parameters to the proteomic and phenotypic measurements in 54 drug combination conditions and at 8 time points. This results in a model containing 124 molecular and two phenotypic nodes. For a complete description of the model equations, see Materials and Methods.

#### Model selection and error estimation

Datasets are subdivided into training, validation and test sets (referring to the gray, green and blue boxes, respectively, in Figure 3 top left). Model parameter estimation is performed on the training dataset. We varied the regularization parameter *λ* on a linear grid and trained 10 network model per grid value. As expected, higher values of *λ* result in sparser networks with fewer interactions (Figure 3 top right, magenta) and higher error on the training data (Figure 3 bottom left, gray). The optimal regularization strength is selected using the validation dataset by (i) Bayesian Information Criterion (BIC) computed on the training set (Figure 3 top right, gray) and (ii) by residual sum of squares (RSS) (Figure 3 bottom left, green). This regularization parameter was determined to be *λ*\* = 3 for both the BIC and RSS metric (Figure 3, bottom left). The expected generalization error is computed using the test dataset (Figure 3 bottom left, green). We studied the accuracy of the final models on the independent test dataset (Figure 3 bottom right). The generalization error for optimal *λ*\* on left-out data was on average 0.17 for each of the 10 models. The Pearson correlation coefficient between model prediction and the experimentally measured data was 0.54 for all data (molecular and phenotypic) and 0.79 for the phenotypic data alone. The correlation is found to be higher when looking only at the later time points of molecular and phenotypic data (Pearson correlation coefficient 0.71 for the last three time points (24, 48, and 67 hours) and 0.74 for the last time point, Figure 13). We conclude that the method for network modeling has a reasonably accurate predictive power, especially for predictions in the 24–67 hour range and that the models are useful for predicting the effects of new perturbations as hypotheses. Reasonable accuracy means that an affordable number of experiments would lead to a positively validated result (a ‘hit’) that can be advanced to pre-clinical investigation.

#### Analysis of network models

We used the optimal regularization parameter *λ*\* = 3 to infer 101 network models on all available data. To have reasonable diversity, the models were inferred without using prior information of known biological interactions. Several key model interactions agreed with interactions reported in the literature. For example, the inferred effect of selected drugs on proteins (Figure 4) is in agreement with known drug–protein interaction patterns (i.e., MEKi inhibits the phorphorylation of ERK1/2 at T202/Y204 and RAFi inhibits MEK1/2 phorphorylation at S217/221) as well as unknown interactions (i.e., PKCi inhibits phorphorylation of CREB at S133). Moreover, these model-derived drug–protein effects are in agreement with the observed drug effects from single-drug RPPA measurements (Figure 8).

**Figure 4:**
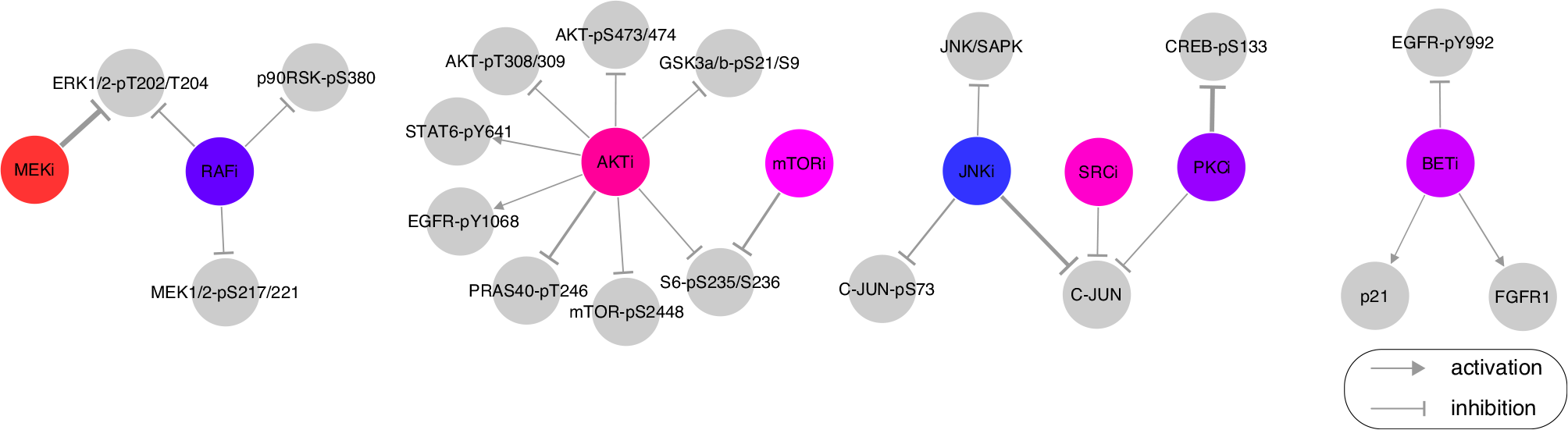
Model-inferred effect of drugs on proteins and phospho-proteins. The effect of drug treatment on (phospho-)protein levels is captured as edges between drugs (colored circles) and proteins (gray circles) in the model. Some of the edges are well known (e.g., MEKi inhibits ERK1/2-pT202/T204) and some appear to be novel or indirect (e.g., inhibitory effect of PKCi on CREB-pS133). Based on the distribution of edge values over the 101 network models, we show only the strongest drug–protein edges for visualization purposes (85th percentile for positive/activating interactions and 15th percentile for negative/inhibiting interactions, by absolute value).

Saturation effects of drugs are taken into account by the model using the open parameter *δ*_*i*_. For values of 0 < *δ*_*i*_ < 1, we have an almost linear drug effect; values of *δ*_*i*_ > 1 do not result in any significant change as drug concentrations increase. For the case of PKC and SRC inhibitors, we find a *δ*_*i*_ in the linear range, i.e., that a doubling of the low dose results in almost the double response, while RAF and JNK inhibitors have almost reached saturation, i.e., no significant difference between the drug effect in low and high dose of the drugs (Figure 9).

#### Predicted effects of drug perturbations and nomination of targets for drug testing

Using the 101 inferred network models, we predicted the phenotypic response (cell growth and apoptosis) as a function of network node inhibition (proteins and phospho-proteins). In particular, we systematically simulated a wide spectrum of inhibition strengths (*c*^pert^) applied to every molecular node (equation 7) and simulated the phenotypic response according to equation 8 at *t* = 72 h (see Materials and Methods). The mean resulting dose response curves for every model when inhibiting each molecular node were used to estimate half maximal effective concentrations (EC_50_) for cell growth and apoptosis. We used these EC_50_ concentrations when simulating the effect of pairwise combination perturbations of the 124 proteins and phospho-proteins on each phenotypic node independently. The predicted responses were averaged across the individual model predictions and ranked. The 20 highest-ranked nodes were further analyzed (Figure 5, left: cell growth reduction, right: apoptosis increase). In total, we predicted cell growth and apoptosis for 124 · 123 = 15,252 conditions for 101 networks, resulting in 1,540,452 simulated responses.

**Figure 5:**
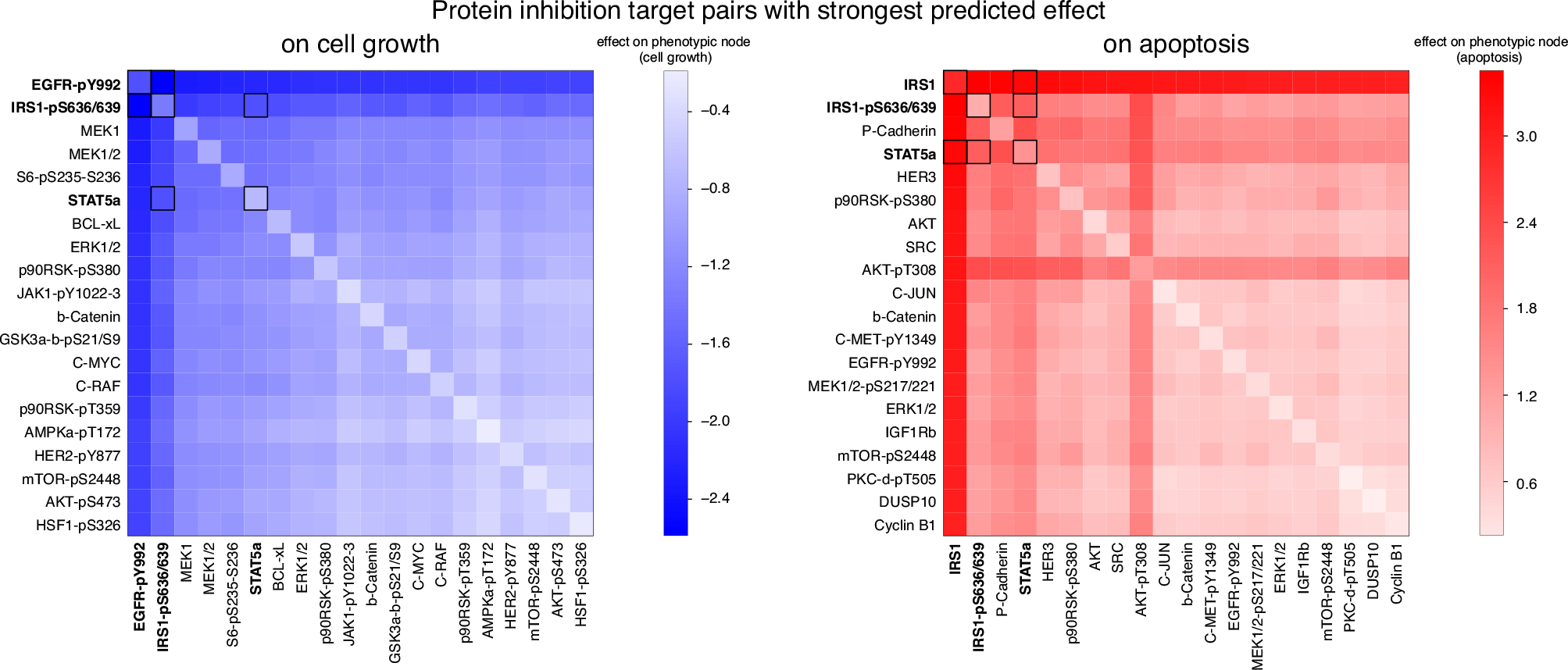
Model predicts the effect of combination perturbations and suggests optimal inhibitor combinations. The top 20 × 20 predictions of pairwise inhibition of molecular nodes (i.e., proteins and phospho-proteins) that decrease cell growth (bottom left, blue) and increase apoptosis (bottom right, red). Cell growth and apoptosis are computed for each target combination as the average over 101 network model predictions (diagonal represents predictions for inhibition of single targets). Combinations nominated for drug testing are highlighted by dark-rimmed squares. (For the predicted values of experimentally tested combinations with an inhibitor of p38-MAPK-pT180/T182 see Figure 6 top left.)

The most effective predicted combination for decreasing cell number was found to be inhibition of the phosphotyrosine at position 992 on EGFR together with phospho-serine at positions 636/639 on IRS1 (Figure 5 left). Simultaneous inhibition of the protein IRS1 together with reduction of phospho-serine at positions 636/639 on IRS1 was found to the most effective combination for increasing apoptosis (Figure 5 right). Note that IRS1 alone had the nearly the same predicted effect on apoptosis as the combination.

EGFR is involved in growth signaling through the RAS/MEK pathway and EGFR-pY992 is an activating auto-phosphorylation site at the receptor [13]. There are many available drugs that specifically inhibit EGFR and are expected to decrease EGFR-pY992 as a proxy of downstream action, for example the drugs lapatinib [14], gefinitib, and erlotinib. Inhibitors of EGFR alone did not have a strong anti-proliferative effect in melanoma cell lines [15].

IRS1 transmits signals from IGFR to PI3K/AKT. The protein can be inhibited by the compound NT157 [16], which has two mechanisms of action: 1) reduce IGF1 signaling through dissociation of IRS1 from the IGF1 receptor, and 2) reduce levels of IRS1 by phosphorylation at serine sites that lead to proteosomal degradation [16]. IRS1-pS636/639 is one such serine phosphorylation site [16]. The inhibitor NT157 is reported to first increase IRS1-pS636/639, which induces degradation of IRS1 (within hours) [16]. It has been reported that NT157 prolonged survival in the RAF inhibitor resistant melanoma cell line A375, both through reduced IGF1-signaling and reduced STAT3-signaling [17]. The inhibitor NT157 has also been shown to increase the tyrosine-phosphorylation 1172 at EGFR in the melanoma cell line A375 [18]. Using an EGFR inhibitor to abrogate this increase in EGFR-pY1172 after NT157 treatment may provide a mechanism for increased drug efficacy. In summary, simultaneous inhibition of IRS1 and EGFR is predicted to both reduce cell growth and increase apoptosis. Its effect can in theory be achieved by using the drugs NT157 (IRS1 inhibitor) and gefitinib (EGFR inhibitor) (Figure 5 bottom middle).

### Experimental test of model predicted drug effects

To test the predictive potential of our model, we selected a subset of the 124 molecular nodes for experimental testing. Nodes were chosen based on i) those that had the strongest predicted effect i.e., EGFR-pY992 and IRS1-pS636/639, ii) those that cover a wide range of predicted response patterns from substantial growth reduction to no effect on growth (Figure 6 top), and iii) those that were targetable by available drugs (Figure 6 top right). Based on these criteria, we selected gefitinib and NT157 to target EGFR-pY992 and IRS1-pS636/639, respectively. The drug CAS285986, which inhibits STAT5b, was chosen as a proxy for STAT5a inhibition, and the p38-MAPK inhibitor SB203580 was selected to target p38-MAPK-pT180/T182. We applied seven different concentrations of each drug, alone and in all pairwise combinations, and measured cell number in three-hour steps from time 0 until 96 h in three replicates (Materials and Methods).

**Figure 6:**
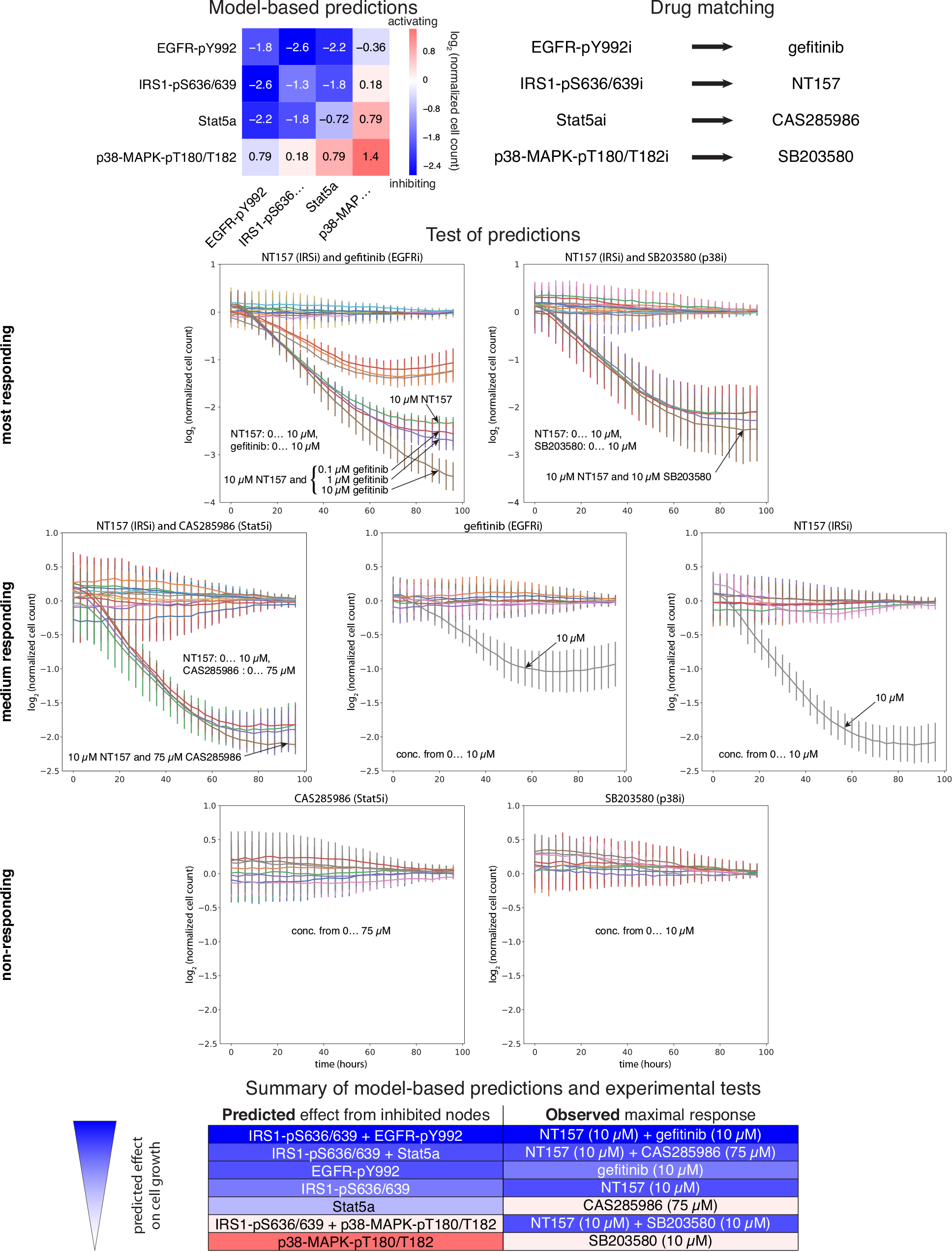
Experimental testing of model predictions using single and pair drugs. Several single and pairwise perturbations that were predicted to have a variety of effects were experimentally tested in the melanoma cell line A2058 (top left; see main text). Cells were perturbed with drugs that target nodes in the computational model (top right). During perturbation, cells were subjected to live-cell imaging, the resulting images were segmented, and cell count was quantified, normalized relative to no-drug control and log_2_-transformed (see growth curves in middle panel). Agreement between model prediction and experiment is summarized (bottom panel).

**Figure 7:**
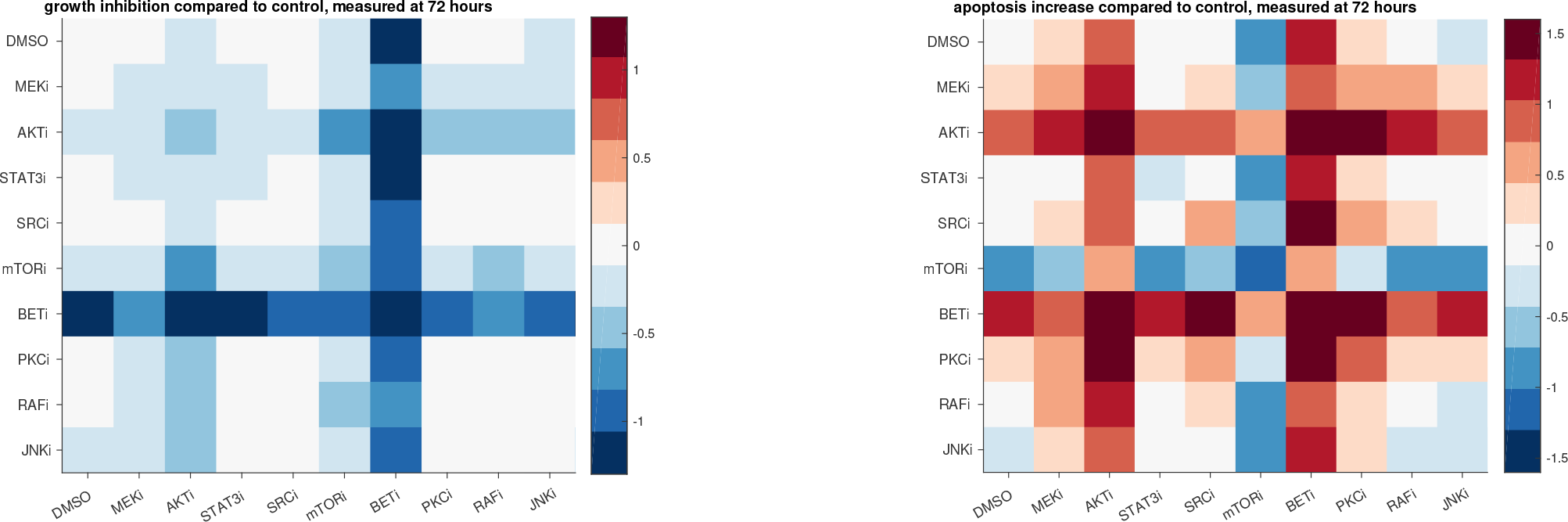
Effect of drugs after 72 hours of drug treatment compared to untreated controls (DMSO). Drug-induced decreased cell growth (left) and apoptosis (right) compared to untreated cells. The response to single drugs in both high (2x low dose) and low dose as well as all double drug combinations in the low dose were measured. The low dose of the drugs are displayed in the top row/left column, and the high dose of the drugs in the diagonal.

**Figure 8:**
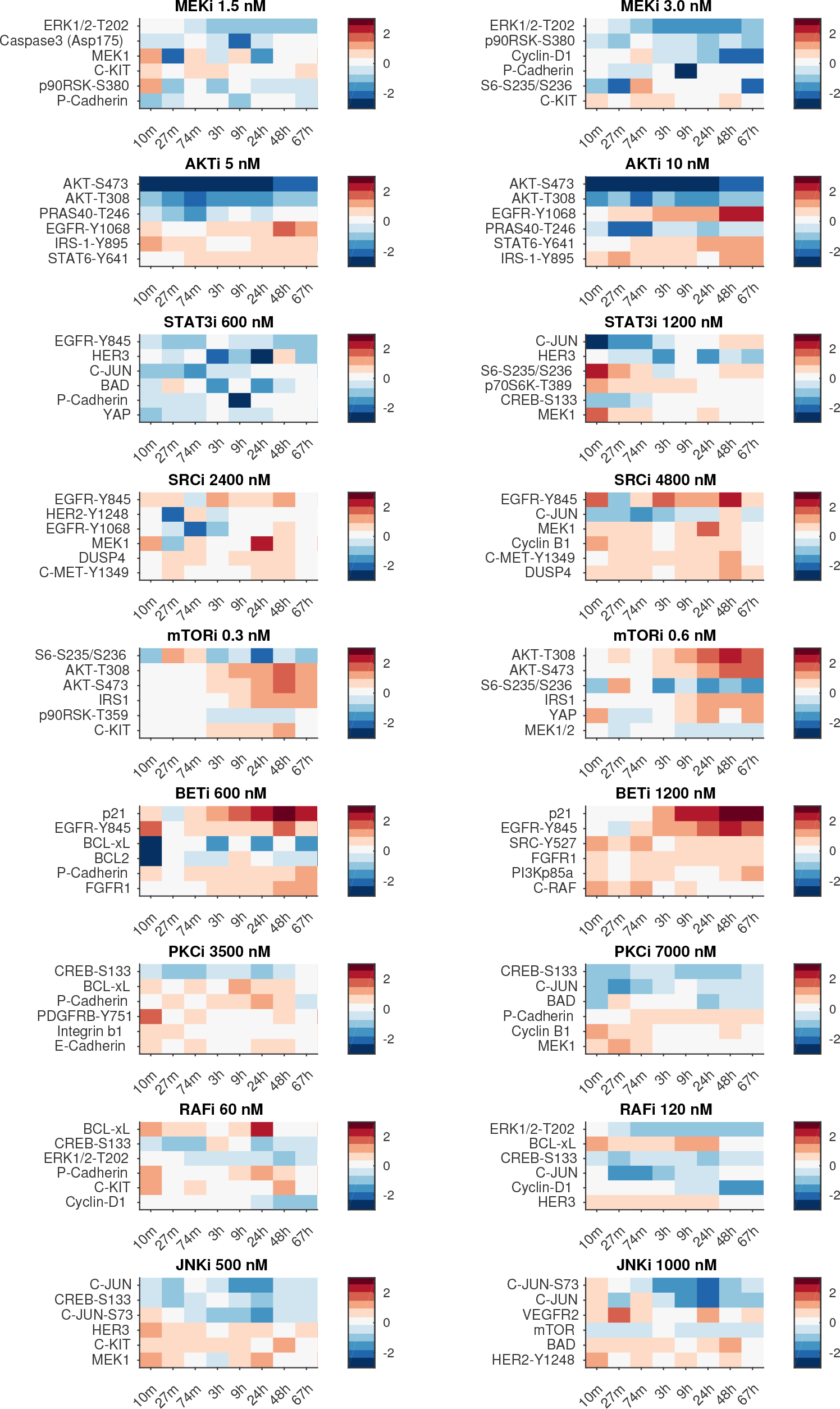
Proteomic drug responses to single drugs at high and low dose: The 6 most responding (phospho-) proteins to single drug perturbations. The data is ranked based on the absolute median response over the time points.

The drug pair with the strongest predicted effect, NT157 and gefitinib, reduced cell growth substantially relative to no-drug control, and the combination is more effective than the effect of either drug alone (Figure 6 middle). The drug pair predicted to have the second strongest effect, NT157 and CAS285986, also reduced cell growth, but to a lesser extent than NT157 with gefitinib, and only for drug combinations that included the highest dose of NT157 (10 *μ*M) (Figure 6 middle, 2nd row). Furthermore, NT157 and gefitinib were both predicted to reduce growth alone, and experimental tests exhibited a reduced cell number with the highest dose tested (10 *μ*M) (Figure 6 middle, 2nd row). CAS285986 alone was predicted to exhibit a slight reduction in cell number, and no effect was observed at any dose of the drug (Figure 6 middle, 3rd row). SB203580 was predicted to activate growth, but no effect of the drug was seen at any concentration (Figure 6 middle, 3rd row). Lastly, although we predicted the combination of NT157 and SB203580 would have little-to-no effect on cell growth (with SB203580 antagonizing the effect of NT157), the experimental data do not support this hypothesis. However, a substantial reduction at the highest dose of NT157 (10 *μ*M) is measured (Figure 6 middle, 1st row). The combination of gefitinib and CAS285986 failed QC and results were inconclusive. In summary, with the exception of the NT157–SB203580 combination, the experimental results largely agreed with the model-based predictions.

## Discussion

We present a method to infer network models from time-resolved molecular and phenotypic perturbation biology data. The perturbation biology data consists of data from multiple drug perturbations, applied as single drugs and drug combinations in a melanoma cell line. The method utilizes the full time-series data, and outputs network models that characterize the interactions between drug perturbations, (phospho-)proteins, and phenotypic changes. The inferred models allow for the prediction of phenotypic responses – cell growth and apoptosis – for unseen perturbations, and can therefore be used to generate large-scale drug discovery hypotheses. As an application, we use the method to predict the most effective drug target combinations in melanoma, and test corresponding drugs experimentally in one case.

When we experimentally test the drug target combination that is predicted to be most effective, i.e., to inhibit the proteins EGFR and IRS1, we find a reduced number of cells only at high doses (10 *μ*M) of the selected drugs. There are several possible reasons for this discordance. The first reason has to do with drug specificity in relation to the proteomic nodes in the network models. The drug gefitinib does not only inhibit the node EGFR-pY992, instead the whole tyrosine kinase domain of EGFR is inhibited. There are several other EGFR-related nodes in our network models. For instance, EGFR-pY845 is predicted to *increase* cell number when reduced (Figure 10). This difference in predicted effect between inhibiting EGFR-pY845 and EGFR-pY992 might hint for a stronger growth reduction at more specific downstream targets of EGFR-pY992. The second reason has to do with antibody cross-reactivity. The EGFR-pY992 antibody can also target HER2, and we might therefore study a combined effect of EGFR and HER2 inhibition. The combined effect is not inhibited by gefitinib since gefitinib is selective for EGFR. There are other drugs that inhibit both EGFR and HER2 (e.g., erlotinib or peletinib) that are more efficient in melanoma cell lines [15]. The third reason has to do with translatability from model predictions to available drugs. No drug is known to directly inhibit IRS1-pS636/639, and, in contrary, one of the mechanism of action of the drug NT157 is to *increase* serine phosphorylations that marks IRS1 for proteosomal degradation [16]. Therefore, even though NT157 decreases IRS1 protein, we have no knowledge on the potential of NT157 to decrease IRS1-pS636/639 significantly. The effect of IRS1 inhibition in cancer has not been fully explored, and there are thus not many drugs to choose from. For these reasons, the model based predicted vulnerability – inhibition of EGFR-pY992 and IRS1/IRS1-pS636/639 in A2058 cells – might be more potent when alternative drugs are chosen or with other methods for inhibition.

**Figure 9:**
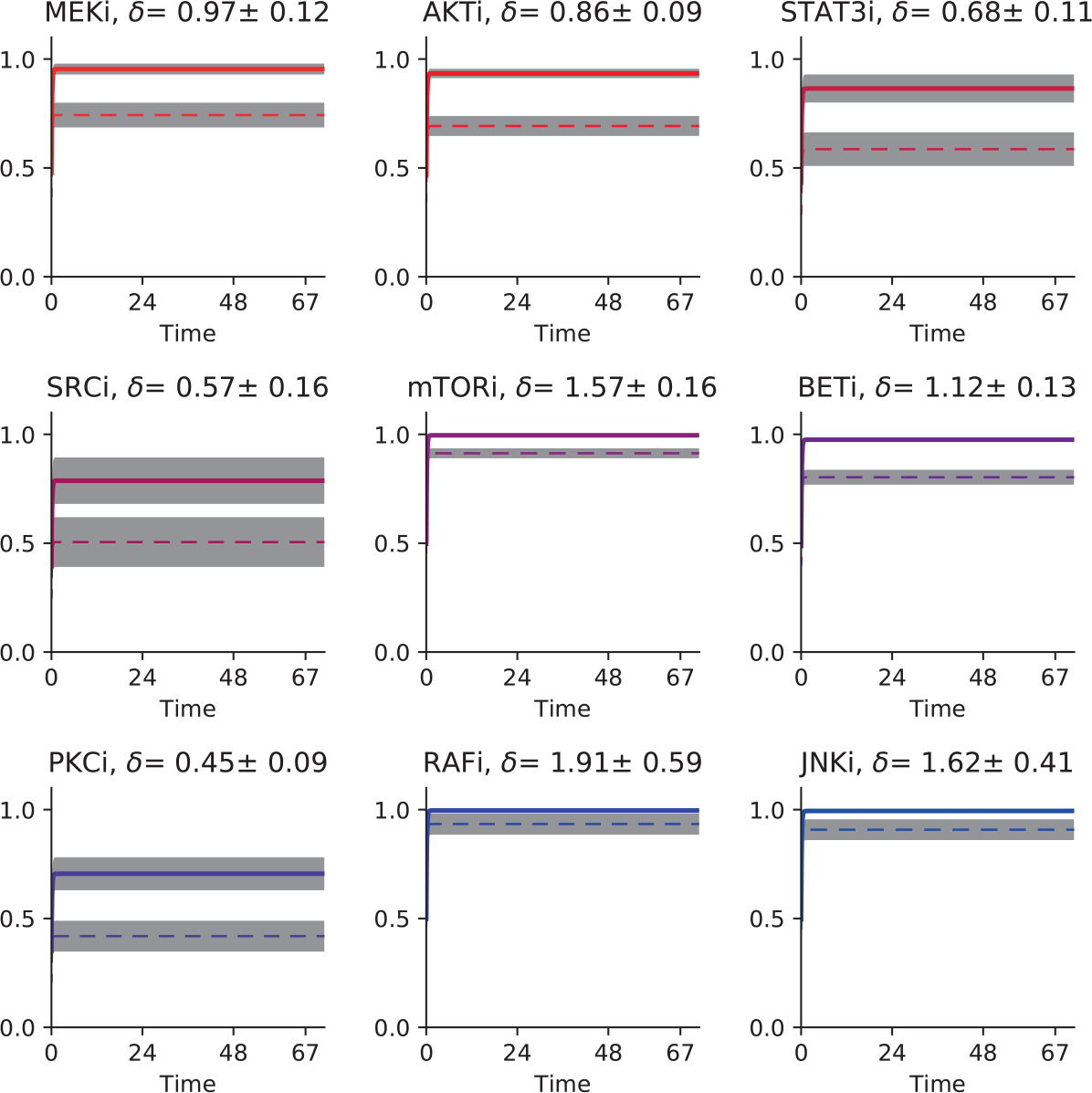
The temporal patterns of the drug nodes. The mean and standard deviation of the simulated drug nodes for the high dose (solid line) and low dose (dashed line) of the drug for the 101 created network models and the corresponding inferred values of *δ*.

**Figure 10:**
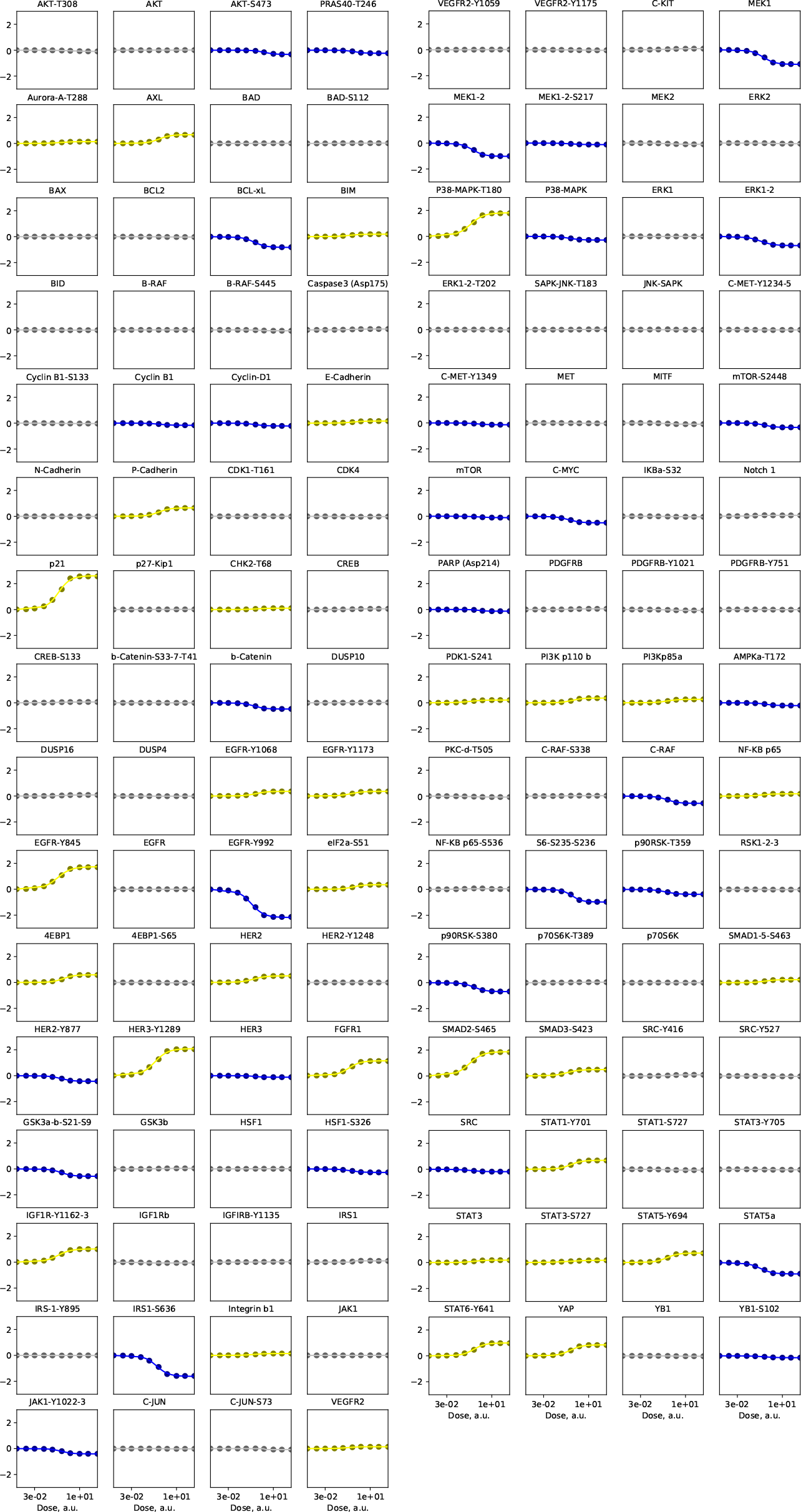
Single node inhibitions, effect on cell growth. Highlighted are the nodes that give at least 2% of the maximal effect. Inhibited nodes that give the desired effect (growth reduction) in blue, and inhibited nodes with the opposite effect (growth increase) in yellow

Our modeling framework is data driven, i.e., network interactions are directly inferred from data without the use of prior knowledge on drug–protein, protein–protein, and protein–phenotype interactions. With this approach, we can subsequently compare the inferred interactions to the literature, and thereby evaluate the quality of the obtained models (see for example Figure 4 for inferred drug–protein interactions). For drug–protein interactions, we believe that this is an important novelty, since many drugs have unspecific or even unknown molecular activity. Even kinase inhibitors that are commonly considered specific and are therefore used to draw conclusions about the physiological roles of molecules, are often not. Indeed, when panels of kinase inhibitors have been profiled against panels of kinases, multiple targets and off target effects have been found in many cases [19–22]. Our framework is useful regardless of the specificity of inhibitors/drugs since all data can be utilized as informative perturbations.

As any data driven modeling method, our developed framework is heavily dependent on both the quality and quantity of data. We have measured the response to 54 drug combinations at 8 time points, and therefore have 432 data points for each node. Even with such a rich data set, we obtain rather big uncertainties in predictions of new drug targets (Figure 12), which hints to the fact that the data is not sufficiently informative to result in more precise predictions. Therefore, to avoid mis-interpretation of predictions, it is important to always study a set of obtained network models, and not only the single best solution.

One limitation of the current approach is the number of nodes that is possible to include, since the optimization problem is quadratic. With the tested 124 (phospho-)protein nodes, each model has around 16,000 parameters that need to be inferred. The use of high-performance compute facilites makes this feasible. Another obvious restriction of the inferred models are their limitation to the available experiments, i.e., only measured (phospho-)protein levels and phenotypes can be included as nodes into the networks. Therefore, interactions between nodes should not be seen as direct biochemical interactions, but logical indirect interactions. For all included nodes, however, we can characterize the response profiles completely, i.e., all combinations of perturbations of included nodes can be tested *in silico*.

The developed framework to infer network models that are constrained by time-resolved responses to perturbations can be applied to cellular systems under the effect of any kind of systematic perturbation, such as CRISPR-Cas9 gene knockouts.

## Materials and Methods

### Genetic features of melanoma cell line A2058

The selection of cell lines that are good models for observed alteration patterns in human tumor tissue is a non-trivial task [23]. In this study, we use the melanoma cell line A2058 for the derivation of optimal drug combinations based on patient-representative genomic profiles [24]. A2058 carries the BRAFV600E mutation that resembles the patient segment which is treatable with the RAF inhibitor vemurafenib and MEK inhibitors such as trametinib. However, A2058 is a RAF/MEK inhibitor resistant cell line with concurrent activation of ERK, PI3K/AKT and cell cycle pathways and TP53 mutation. Specifically, it carries BRAFV600E, MAP2K1P124S and TP53V274F mutations and is altered in tumor suppressor genes like CDKN2A, RB and PTEN [25]. As in most melanoma samples, the cell line A2058 carries a large number of additional genomic alterations other than those listed here [24, 26–28]. For A2058 cells, a synergistic effect of drug combinations has been shown using the combination of BRAF and PI3K inhibition (drugs: PLX4720 and GDC0941, respectively) [26].

### Model equations

For the molecular response model, molecular nodes represent concentrations of proteins and phospho-proteins. We start with Equation 1. Applied perturbations only affect molecular nodes. All interactions between molecular nodes are directional. Self-interaction terms are excluded from the model formulation. The resulting model of the temporal dynamics of the *i*-th **molecular node** *i* = 1, …, *N*_molec_ in experimental condition *μ* is given by,

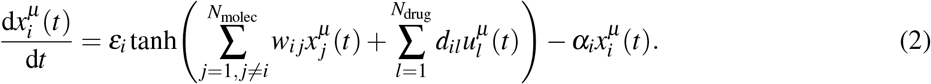

In particular, 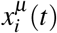 denotes the temporal log_2_-control-normalized protein level (see defining equation in Data normalization below), which is subjected to drug perturbations as defined by the superposition of drug nodes 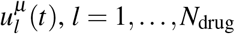 (see below). In order to model saturation effects and cap the time derivatives, we use the hyperbolic tangent as the sigmoidal transfer function. By doing so, the interaction-driven differential change of variable *i* is bounded to values in [−*ε*_*i*_, *ε*_*i*_]. The interaction parameters *w*_*i j*_ characterize the effect of protein level *j* on *i* and *α*_*i*_ describes how quickly the *i*-th protein level returns to the zero-level steady-state in the absence of any interaction or drug. The effect of the drug *l* on molecular node *i* is quantified by *d*_*il*_. In contrast to our previous perturbation modeling approaches [1, 12] in which the interaction between drug node and target were defined based on prior knowledge, in this work we determine the drug–molecular node interactions *d*_*il*_ as a result of model inference. In addition to *d*_*il*_, the parameters *α_i_*, *ε*_*i*_ and *w*_*i j*_ are also data-derived.

Using Equation 1, we define the *l*-th **drug node**, *l* = 1, …, *N*_drug_, by,

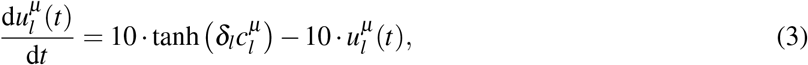

where we enforce a fast-acting drug effect by choosing *ε* = *α* = 10 in Equation 1. Here 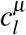 describes the drug concentration relative to the single-drug concentration (i.e., 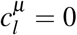 when the drug not present, 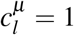 when drug is present at low dose concentration and 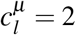 when the drug is present at high dose (i.e., 2× low dose) in experimental condition *μ*. Equation 3 has the analytical solution,

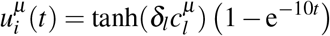

and *δ*_*l*_ is the effective impact, which is inferred in the parameter inference process.

We also use Equation 2 to define the two **phenotypic nodes** cell growth (cg) and apoptosis (ap). These nodes are modeled in the same way as molecular nodes except that interactions are only unidirectional (i.e., from molecular to phenotypic node). These dynamics are described as,

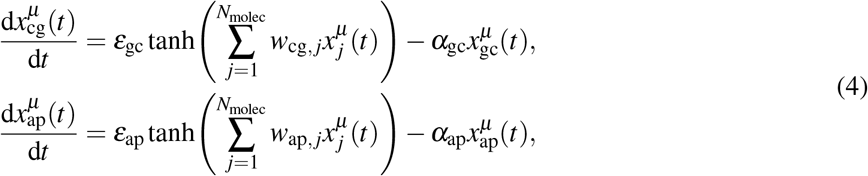

where 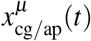 denotes the temporal log_2_-control-normalized apoptotic readout and cell number, respectively (see defining equation in Data normalization below).

### Definition of loss function

The residual sum of squares for a given parameter set Θ, RSS(Θ), is used as measure of agreement between model simulations and observations,

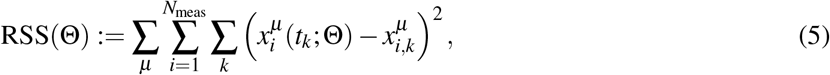

where all free parameters are grouped into 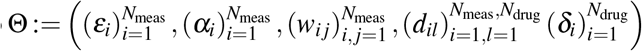 and *N*_meas_ = *N*_molec_ + *N*_phen_ = *N*_molec_ + 2 is the number of measured observables, i.e., the number of proteins and phospho-proteins (molecular nodes), and the two phenotypes. Moreover, 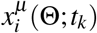 represents the simulated response given the parameters Θ, *μ* is the experimental condition and *t*_*k*_ is the *k*-th time point. The corresponding observed data is denoted by 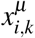. In addition to the minimization, regularization was chosen to reduce the number of edges. In practice, an *ℓ*^1^-norm on the interaction parameters *w*_*i j*_ was added to the problem of minimizing the residual sum of squares as function of Θ

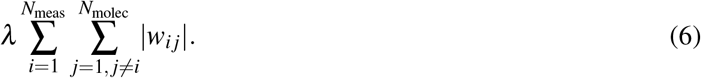

where *λ* sets the regularization strength and was chosen based on data holdout and Bayesian Information Criterion considerations.

The loss function was minimized using Adam, an implementation of a stochastic algorithm for first-order gradient-based optimization [29]. We used the version of Adam included in TensorFlow (Google Brain team, Mountain View, CA, U.S.A.), and the entire simulation and optimization procedure was performed in a python environment. For all minimization runs, we used a fixed learning rate of 0.01 and evaluated the results when 10,000 iterations or 24 hour run time were reached, whichever occurred first. Failed optimization runs with all *w*_*i j*_ = 0 were discarded.

The Adam minimizer does not offer soft thresholding, so no exact zeros of parameters are obtained even when the *ℓ*^1^-norm is applied. We adopt an alternative strategy to introduce parameter sparsity by cropping the interaction parameter *w*_*i j*_ in each update step *n* ↦ *n* + 1, for interactions from protein/phenotypic nodes: 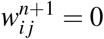 if 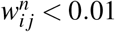, and for interactions from drugs nodes: 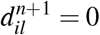 if 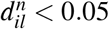.

### Model selection

We adopted two strategies to select the most informative model without overfitting to the given data using the Bayesian information criterion and (2) by division of the input data into training, validation, and test datasets.

#### Bayesian information criterion

Assuming independent and identically distributed model errors drawn from a normal distribution, the maximum loglikelihood 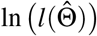 can be expressed in terms of the minimum RSS 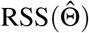, where 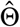 is the parameter set that maximizes the likelihood function. We use the following simplified definition of the Bayesian information criterion, BIC,

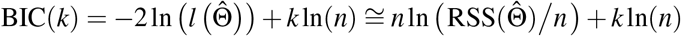

where *k* is the number of non-zero model parameters and *n* is the number of data points used in the parameter estimation. The number of non-zero parameters is a function of the regularization applied via *λ*, i.e., *k* = *k*(*λ*). We select the optimal regularization parameter *λ* as the minimum of the BIC,

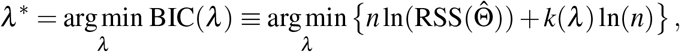

where we omitted contributions from constants that are dependent on the number of data points *n*. For the current data set, the optimal regularization parameter is found as *λ*\* = 3 for the Bayesian information criterion (Figure 3 top right). The Akaike information criterion did not have a clear minimum.

#### Dataset division

In addition to selecting *λ* using the Bayesian Information Criterion, the optimal regularization parameter *λ*\* is evaluated by splitting the data into a training dataset to estimate the model parameters, a validation dataset to determine an optimal *λ*, and a test dataset to evaluate model performance. The training set is chosen to contain all single drug conditions, and the test sets are chosen as a split between all combinations of drugs (Figure 3 top left). As with the Bayesian Information Criterion, the optimal regularization parameter is found to be *λ*\* = 3 (Figure 3 bottom left).

### Systematic *in silico* perturbation of molecular nodes

In order to identify new potential drug target combinations, we combine simulations using models built with all the data with an *in silico* inhibitor of each of molecular node, 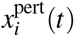. The *in silico* inhibitor of molecular node *i* is added to the model equations as a step concentration 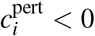 for *t* ≥ 0 and then we simulate the protein and phospho-protein dynamics to the last measured time point *t* = 72 h. The resulting dynamics for the perturbed/inhibited molecular node *i* = 1, … *N*molec are found from,

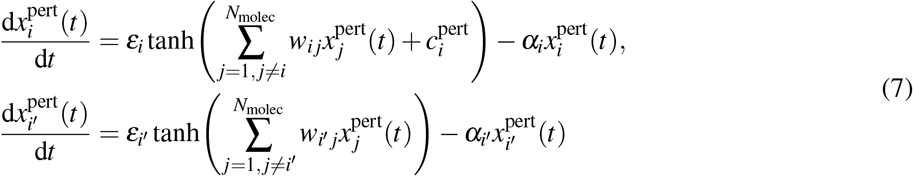

with *i*′ = 1, …, *i*− 1, *i* + 1, …, *N*molec. The response of the phenotypic nodes, 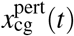 and 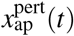, are determined as,

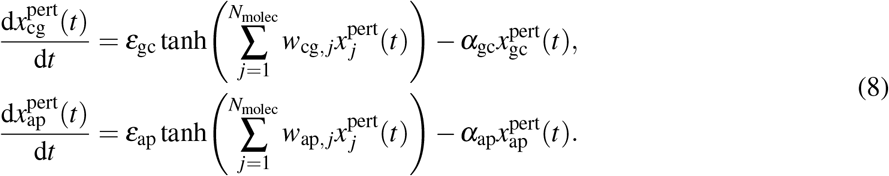

#### Selection of inhibition strength for *in silico* combination perturbations

We assume that at *t* = 72 h for a single *in silico* perturbation of molecular node *i* with concentration 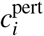 that phenotypes as the solution to Equations 7 and 8 follow the shape,

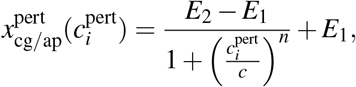

where 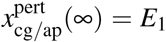 and 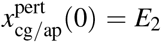 represent the maximum and minimum change in each phenotype, respectively. 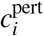 is the concentration of the *in silico* inhibitor specifically targeting molecular node *i*. *n* and *c* are open parameters. *n* is the Hill coefficient and *c* is the *in silico*-derived 50%-effect concentration, EC_50_. In our framework, we focus on the effect on cell growth and apoptosis by single molecular node inhibition. As the data are all log_2_-normalized to control data, 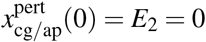 and so the equation above simplifies to,

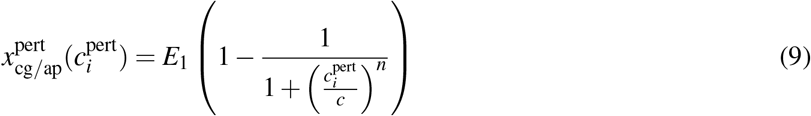

with 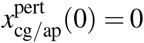 and 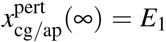 (see Figures 10 and 11 for individual dose response of apoptosis and cell growth as a function of inhibiting single molecular nodes). These *in silico*-derived EC_50_ are subsequently used for molecular node inhibition simulations. These simulations are systematically carried out such that all single and pairwise molecular nodes are perturbed, and their effect on cell growth and apoptosis is calculated (Figure 5).

The scale of concentration in the simulations, and hence the estimated *in silico* EC_50_ concentration (parameter *c* in Equation (9), Figure 10), is expressed in arbitrary units, but can nevertheless be related to the experimental EC_50_ concentration. This can be done by equivalencing the estimated *in silico* and experimentally determined EC_50_ values. This assumes however that the inhibitor used in the experiments is of perfect specificity.

**Figure 11:**
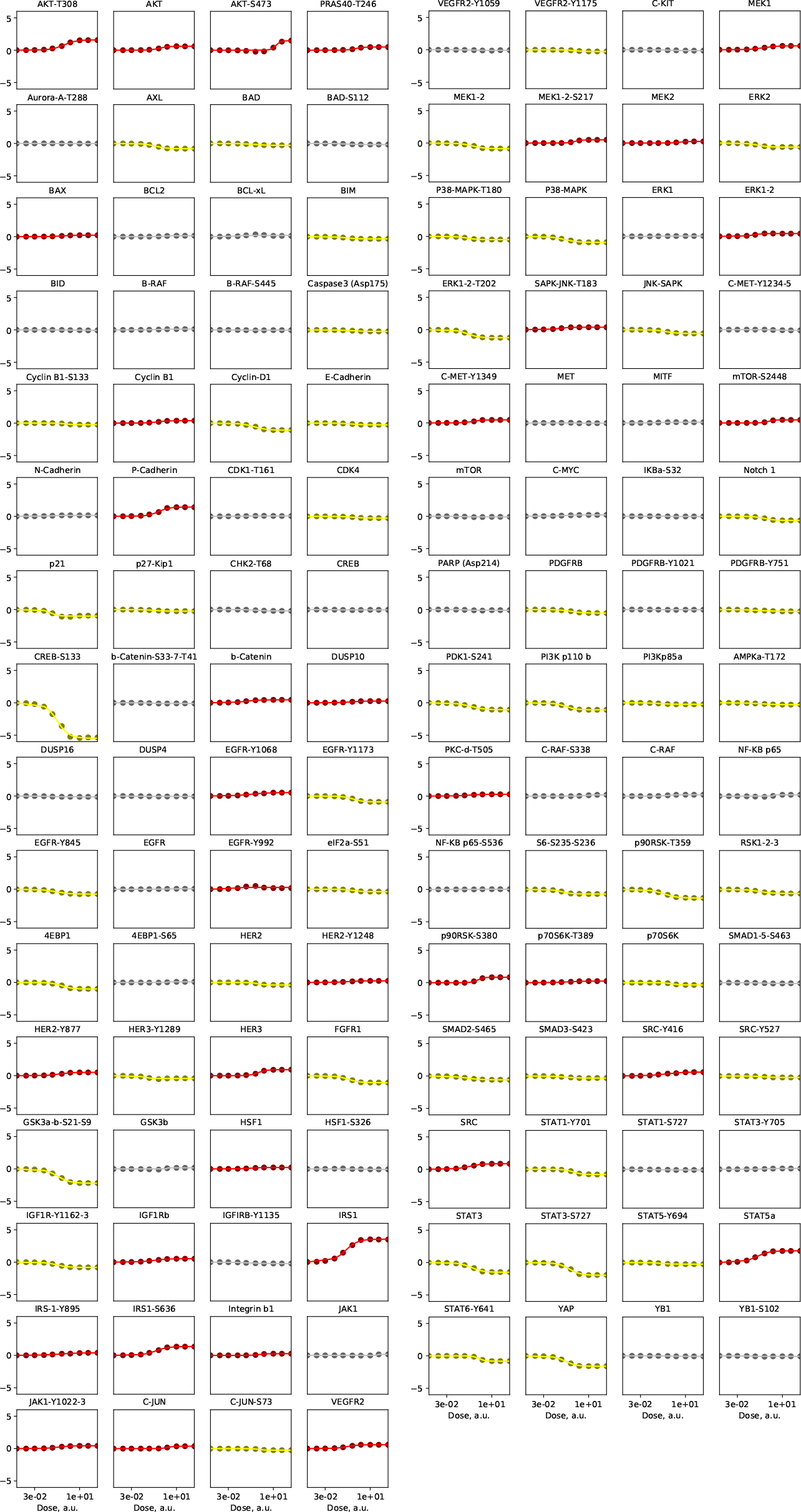
Single node inhibitions, effect on apoptosis. Highlighted are the nodes that give at least 2% of the maximal effect. Inhibited nodes that give the desired effect (apoptosis increase) in red, and inhibited nodes with the opposite effect (apoptosis reduction) in yellow

**Figure 12:**
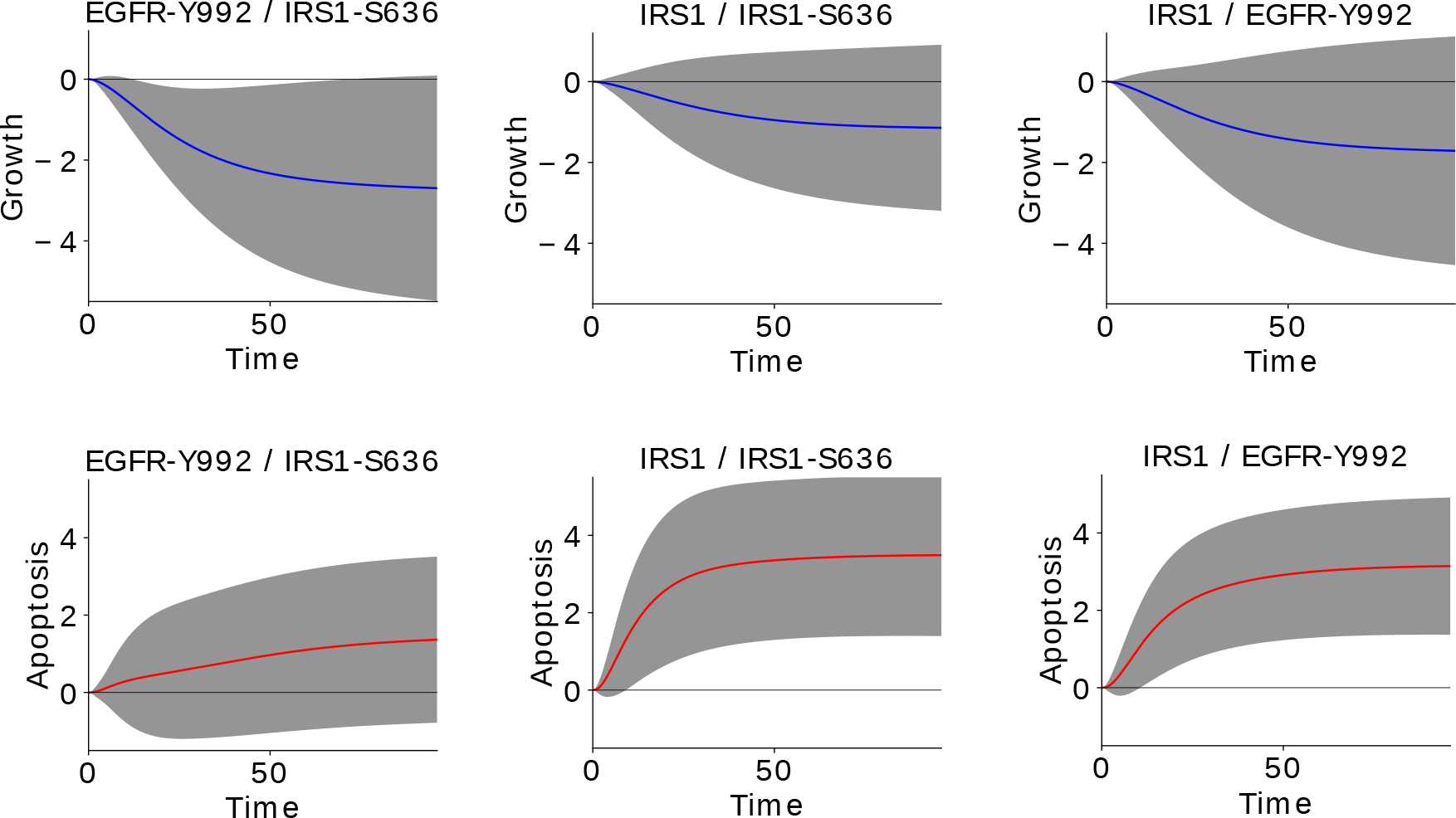
Predicted temporal patterns of growth and apoptosis in response to pairwise combination perturbations (EGFR-pY992/IRS1, EGFR-pY992/IRS1-pS636/639, and IRS1/IRS1-pS636/639). The mean (yellow line) and standard deviation (gray area) of simulations of the 101 created network models.

**Figure 13:**
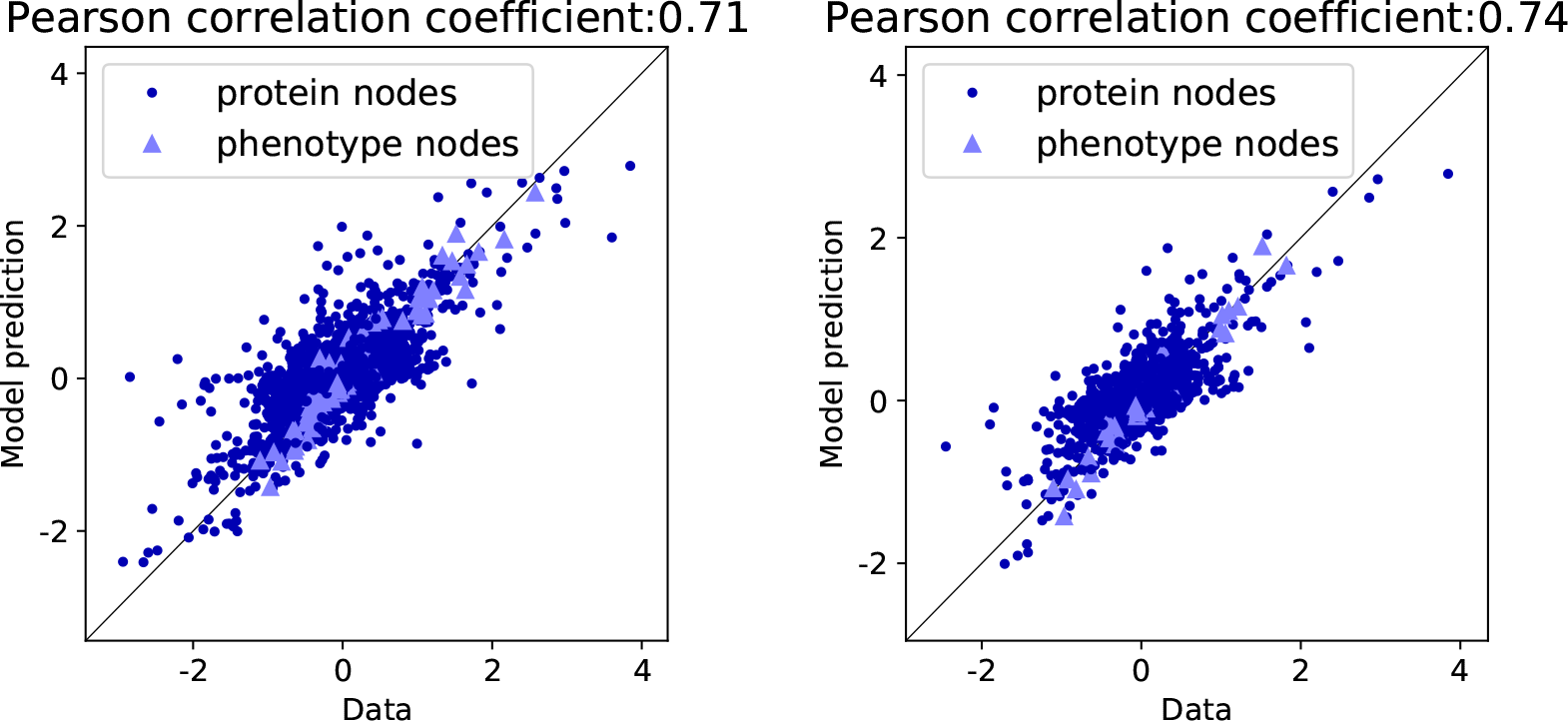
The correlation between model simulations and data is higher for later time points. Comparison between prediction and data for the last three measured time points: 24, 48, and 67 hours (left panel) and the last time point: 67 hours alone (right panel). This result suggests that the model predictions are less reliable in earlier time points. This is probably due to the transitory nature of the drug response and the experimental noise at earlier time points in the data. Related to Figure 3.

## Experimental and data normalization protocols

### Molecular measurements in response to perturbation

The melanoma cell line A2058 was seeded in 6-well plates at 50,000 cells per well. Each biological replicate was run on separate occasions. The physical difficulties and logistical complexity of the experiments required that each time point be run in batches with the early time points being harvested immediately and the late time points being setup immediately following time point harvesting. Perturbed cells were lysed in CLB1 buffer (Bayer Technology Services, Leverkusen, Germany; now NMI TT Pharmaservices, Reutlingen, Germany). For Dataset 1, we performed a 4-fold dilution series of the samples using the Biomek FXP Laboratory Automation Workstation (Beckman Coulter Inc., Brea, CA, U.S.A.) automatic pipetting system in one technical replicate [10]. For Dataset 2, [a single lysate sample was printed twice (technical replicate)] each lysate sample was printed at the respective protein concentration in two technical replicates. Diluted samples were printed onto tantalum pentoxide-coated glass chips and then blocked with an aerosol BSA solution. The protein array chips are then washed in double-distilled H_2_O and dried before measurement. For the immunoassay, we incubated the chips with primary antibodies for 24 hours followed by 2.5 hours incubation with Alexa Fluor-647 conjugated secondary antibody detection reagents. Antibodies are diluted in CAB1 buffer (Bayer Technology Services, Leverkusen, Germany). The immuno-stained chips were imaged using the ZeptoREADER (Zeptosens/Bayer, Witterswil, Switzerland). The ZeptoView 3.1 software (Zeptosens/Bayer, Witterswil, Switzerland) was used to output the reference net spot fluorescence intensity, RNFI. Included standard global and local normalization of sample signal to the reference BSA grid is used.

#### Sample lysis and RPPA chip layout and analysis

All experimental perturbations and sample preparations were carried out in the Sander lab. Cell lysate was aliquoted into multiple samples, and RPPA measurement using the zeptosens platform was carried out in both the Sander lab (MSKCC, Dataset 1) and the Pawlak lab (NMI TT Pharmaservices, Dataset 2).

##### Differences between each dataset

Dataset 1 consists of 71 antibody measurements of three biological replicates at eight time points (10 min, 27 min, 72 min, 3 h, 9 h, 24 h, 48 h, 67 h). For Dataset 1, each RFI data point, 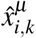, was derived from a four-spot protein dilution series of 0.2, 0.15, 0.1, and 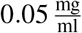. Dataset 2 consists of 86 antibody measurements of three biological replicates at (a) eight time points for biological replicate 1 and (b) two time points (48 h, 67 h) for biological replicate 2 and 3. For Dataset 2, protein lysate was diluted to 3 *μ*g/*μ*l, when concentration was higher than 3.4 *μ*g/*μ*l, and each RFI data point was derived from the mean of two measured spots. Median-centered protein factors, determined for each sample by NMI TT Pharmaservices, were used to normalize protein content.

##### Similarities between each dataset

All data were acquired from the same RPPA platform (Zeptosens/Bayer, Witterswil, Switzerland). 33 antibodies were measured in both datasets as controls as controls, which resulted in 71 + 86 − 33 = 124 unique antibodies. After outlier detection and loading normalization (see above), the median of the three biological replicates from Dataset 1 and the median of the data (one biological replicate in six time points and three biological replicates in two time points) from Dataset 2 was combined and used in the downstream analysis and modeling steps.

#### Fluorescence signal evaluation

Each spot measured on the Zeptosens array corresponds to the relative concentration of a specific protein or phospho-protein in a single condition. This value is detected by measuring the secondary antibody fluorescence that is bound to the (phospho)protein-specific antibody. The relationship between this signal and the absolute concentration of the (phospho-)protein is assumed to be linear in the observed range, i.e.,

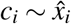

given constant exposure times across all the experimental conditions and time points in each antibody measurement. In Dataset 1, for each (phospho-)protein *i* one measures four antibody fluorescence signals 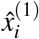, 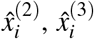 and 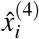 corresponding to the four serial dilutions of cell lysate 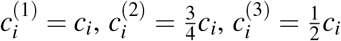 and 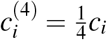 These values are referred to as Referenced Net spot Fluorescence Intensity (RNFI). This allows to quantify the degree of “non-linearity” of the signal by computing the discrepancy between linear fit and actual signal. Assuming a linear dependency, then *x̂*_*i*_(*c*) = *β*_0_ + *β*_1_*c*, the fitting parameters are found in a least-squares estimation,

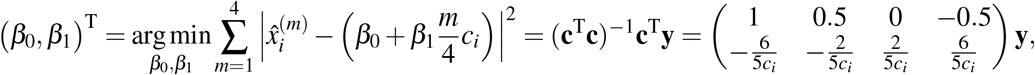

where,

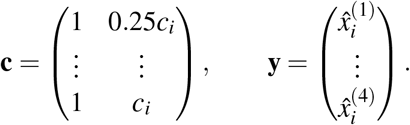

For ideal profiles, i.e., without intercept or *x̂*_*i*_(0) = *β*_0_ ≈ 0, we have *x̂*_*i*_(*c*_*i*_) ≈ *β* _1_*c*_*i*_ (and consequently, 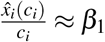). A single value, the Referenced sample Fluorescence Intensity (RFI), corresponding to the linear fit of the 4-spot array corresponding to the mean concentration value, *c* = 0.625*c̄*, is chosen,

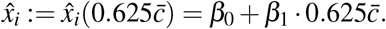

Zeptosens calls the RFI using the reference concentration of the total protein content per spot 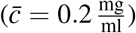. This concentration was chosen to be roughly the protein content of a eukaryote cell. Ideally, the concentrations for each sample are *c*_*i*_ ≈ *c̄*, but in practice *c*_*i*_ varies due to experimental uncertainties. We could estimate *a posteriori* the actual protein loading quantification with the reference loading chip and subsequently adjust *x̂*_*i*_ for each sample by setting *c̄* = median_*i*_{*c*_*i*_}. For Dataset 2, we adopt this strategy and use on-chip measured median-centered protein factors determined from protein stain assays. However, for Dataset 1 we had no measurements of the protein loading and therefore applied double-median normalization (see [27]) for protein loading normalization.

#### Spotting error detection

For Dataset 1, due to stochastic sampling, low concentrations or saturated sample, as well as occasional robotic errors, some lysate spots in the serial dilutions do not follow the expected linear and monotonically decreasing dilution profile. We identify outliers that deviate strongly from the linear interpolation of the RNFI values using the Cook’s distance [30]. We then use a Cook’s distance of 0.8 as cutoff. For Dataset 2, the spotting quality appeared to be more consistent and no spotting error removal was required.

#### Protein loading normalization

Occasionally, samples have total protein concentrations that deviate from the optimal loading concentration. To normalize for this uneven loading in the we perform double-median normalization (sometimes referred to as “loading control”) as described in [31]. For Dataset 1, the log_2_-transformed and normalized value for the *i*-th antibody in time-point *k* and under experimental condition *μ* is then calculated as,

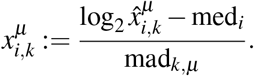

Here, 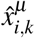 denotes the inear raw data of antibody *i* measured in time point *k* under experimental condition *μ*, 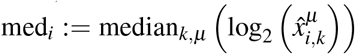 the median of the log_2_-transformed values in each antibody across time points and experimental conditions and 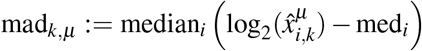 its median absolute deviation. In Dataset 2, in contrast to Dataset 1, we have an estimate on the relative deviation from the target protein loading concentration in each sample of 0.3 *μ*g/*μ*l. The observed protein loading was lower and found to have a median concentration of 0.22 *μ*g/*μ*l). Based on these measurements, we normalize each lysate sample by the corresponding on-chip measured median-centered protein factors.

#### Data normalization

Before applying any further modeling and inference, we subtract the log_2_-transformed and double-median normalized RPPA data in the DMSO control condition (without perturbation) from all previously log_2_-transformed and double-median normalized RPPA data, 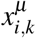:

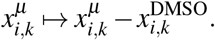

### Phenotypic measurements in response to perturbation

At the start of the experiment, 1,000 A2058 cells were seeded in three biological replicates in 96-well plates. After 24 hours, drugs were added to each well and cells were subjected to real-time imaging using the IncuCyte (Essen BioScience, Ann Arbor, MI, U.S.A.) live-cell imaging system. Images were taken every 3 hours in 3 channels (phase, GFP, RFP) at 4 regions on the plate (12 images per well at each time point). The RFP channel was used to detect nuclear-localized mCherry (constitutively expressed). To detect apoptosis, a Caspase-3/7 green reagent (Essen BioScience) was added to each well. Image analysis using the Incucyte software was used to segment and count all cells (RFP channel) and cells going through apoptosis (GFP channel) at each time point. These counts were log_2_-transformed and normalized relative to DMSO control condition.

### Cell number measurements of optimal drug combinations

#### Cell line

Cell line A2058 with H2B-mCherry was thawed, tested for mycoplasma, and passaged twice prior to testing. The cell line was passaged fewer than 15 times and screened with the compounds within a month of thawing. The cell line was passaged and cultured on DMEM media with 10 % FBS and 1% Penicillin/Streptomycin.

#### Screening Protocol

Cells were harvested, counted, and deposited into 384 well plates at a total volume of 50 *μ*l per well with a *t*_0_ seeding density of 750 cells/well using a Thermo Multidrop Combi dispenser (Thermo Fisher Scientific, Waltham, MA, U.S.A.). Cells were immediately placed into the Incucyte microscope culture and allowed to attach and proliferate for 24 hours. Four compounds (NT157, CAS285986, gefitinib, SB203580) were dissolved into 10 mM solutions two hours prior to dosing. The compounds were dispensed into the previously seeded 384 well plates using a HP D300e Digital Dispenser 24H after cell seeding in a dose matrix testing each concentration point of each drug in combination with each concentration point of the other drugs, with three of the drugs (NT157, gefitinib, and SB203580) titrated from 0.01 *μ*M to 10 *μ*M and CAS285986 titrated from *μ*M to 75 *μ*M. The plates were then placed back into the Incucyte and cell measurements were recorded every 3 hours for 96 hours.

## References

[1] Anil Korkut, Weiqing Wang, Emek Demir, Bulent Arman Aksoy, Xiaohong Jing, Evan J Molinelli, Ozgun Babur, Debra L Bemis, Selcuk Onur Sumer, David B Solit, Christine A Pratilas, and Chris Sander. Perturbation biology nominates upstream-downstream drug combinations in RAF inhibitor resistant melanoma cells. eLife, 4:e04640, 2015.

[2] Richard G. Abramson. Overview of Targeted Therapies for Cancer, 2017.

[3] Paul B. Chapman, Axel Hauschild, Caroline Robert, John B. Haanen, Paolo Ascierto, James Larkin, Reinhard Dummer, Claus Garbe, Alessandro Testori, Michele Maio, David Hogg, Paul Lorigan, Celeste Lebbe, Thomas Jouary, Dirk Schadendorf, Antoni Ribas, Steven J. O’Day, Jeffrey A. Sosman, John M. Kirkwood, Alexander M.M. Eggermont, Brigitte Dreno, Keith Nolop, Jiang Li, Betty Nelson, Jeannie Hou, Richard J. Lee, Keith T. Flaherty, and Grant A. McArthur. Improved Survival with Vemurafenib in Melanoma with BRAF V600E Mutation. New England Journal of Medicine, 364(26):2507–2516, 6 2011.

[4] Jose Luis Manzano, Laura Layos, Cristina Buges, Maria de Los Llanos Gil, Laia Vila, Eva Martinez-Balibrea, and Anna Martinez-Cardus. Resistant mechanisms to BRAF inhibitors in melanoma. Annals of translational medicine, 4(12):237, 6 2016.

[5] Clinicaltrial.gov. https://clinicaltrials.gov/ct2/show/NCT03266159. Accessed: 2018-09-06.

[6] Martin L. Miller, Evan J. Molinelli, Jayasree S. Nair, Tahir Sheikh, Rita Samy, Xiaohong Jing, Qin He, Anil Korkut, Aimee M. Crago, Samuel Singer, Gary K. Schwartz, and Chris Sander. Drug Synergy Screen and Network Modeling in Dedifferentiated Liposarcoma Identifies CDK4 and IGF1R as Synergistic Drug Targets. Science Signaling, 6(294):ra85–ra85, 2013.

[7] Eduardo Sontag, Anatoly Kiyatkin, and Boris N Kholodenko. Inferring dynamic architecture of cellular networks using time series of gene expression, protein and metabolite data. Bioinformatics, 20(12):1877–1886, 2004.

[8] Eduardo D Sontag. Network reconstruction based on steady-state data. Essays in Biochemistry, 45:161–176, 2008.

[9] Lise Boussemart, Helene Malka-Mahieu, Isabelle Girault, Delphine Allard, Oskar Hemmingsson, Gorana Tomasic, Marina Thomas, Christine Basmadjian, Nigel Ribeiro, Frederic Thuaud, Christina Mateus, Emilie Routier, Nyam Kamsu-Kom, Sandrine Agoussi, Alexander M Eggermont, Laurent Desaubry, Caroline Robert, and Stephan Vagner. eIF4F is a nexus of resistance to anti-BRAF and anti-MEK cancer therapies. Nature, 513, 2014.

[10] Xiaohong Jing, Weiqing Wang, Nicholas P Gauthier, Poorvi Kaushik, Alex Root, Richard R Stein, Anil Korkut, and Chris Sander. Protein profiling in cancer cell lines and tumor tissue using reverse phase protein arrays. bioRxiv, page 144535, 2017.

[11] Sven Nelander, Weiqing Wang, Björn Nilsson, Qing-Bai She, Christine Pratilas, Neal Rosen, Peter Gennemark, and Chris Sander. Models from experiments: combinatorial drug perturbations of cancer cells. Molecular Systems Biology, 4(1):216, 2008.

[12] Evan J Molinelli, Anil Korkut, Weiqing Wang, Martin L Miller, Nicholas P Gauthier, Xiaohong Jing, Poorvi Kaushik, Qin He, Gordon Mills, David B Solit, et al. Perturbation biology: inferring signaling networks in cellular systems. PLoS Computional Biology, 9(12):e1003290, 2013.

[13] J. Downward, P. Parker, and M. D. Waterfield. Autophosphorylation sites on the epidermal growth factor receptor. Nature, 311(5985):483–485, oct 1984.

[14] Wenle Xia, Robert J Mullin, Barry R Keith, Lei-Hua Liu, Hong Ma, David W Rusnak, Gary Owens, Krystal J Alligood, and Neil L Spector. Anti-tumor activity of GW572016: a dual tyrosine kinase inhibitor blocks EGF activation of EGFR/erbB2 and downstream Erk1/2 and AKT pathways. Oncogene, 21(41):6255–6263, sep 2002.

[15] Tamás Garay, Eszter Molnár, Éva Juhász, Viktória László, Tamás Barbai, Judit Dobos, Karin Schelch, Christine Pirker, Michael Grusch, Walter Berger, József Tímár, and Balázs Hegeds. Sensitivity of Melanoma Cells to EGFR and FGFR Activation but Not Inhibition is Influenced by Oncogenic BRAF and NRAS Mutations. Pathology & Oncology Research, 21(4):957–968, sep 2015.

[16] Hadas Reuveni, Efrat Flashner-Abramson, Lilach Steiner, Kirill Makedonski, Renduo Song, Alexei Shir, Meenhard Herlyn, Menashe Bar-Eli, and Alexander Levitzki. Therapeutic destruction of insulin receptor substrates for cancer treatment. Cancer research, 73(14):4383–94, jul 2013.

[17] E Flashner-Abramson, S Klein, G Mullin, E Shoshan, R Song, A Shir, Y Langut, M Bar-Eli, H Reuveni, and A Levitzki. Targeting melanoma with NT157 by blocking Stat3 and IGF1R signaling. Oncogene, 35(20):2675–2680, may 2016.

[18] Shih-Ping Su, Efrat Flashner-Abramson, Shoshana Klein, Mor Gal, Rachel S Lee, Jianmin Wu, Alexander Levitzki, and Roger J Daly. Impact of the anticancer drug nt157 on tyrosine kinase signaling networks. Molecular cancer therapeutics, 2018.

[19] Susan Klaeger, Stephanie Heinzlmeir, Mathias Wilhelm, Harald Polzer, Binje Vick, Paul-Albert Koenig, Maria Reinecke, Benjamin Ruprecht, Svenja Petzoldt, Chen Meng, Jana Zecha, Katrin Reiter, Huichao Qiao, Dominic Helm, Heiner Koch, Melanie Schoof, Giulia Canevari, Elena Casale, Stefania Re Depaolini, Annette Feuchtinger, Zhixiang Wu, Tobias Schmidt, Lars Rueckert, Wilhelm Becker, Jan Huenges, Anne-Kathrin Garz, Bjoern-Oliver Gohlke, Daniel Paul Zolg, Gian Kayser, Tonu Vooder, Robert Preissner, Hannes Hahne, Neeme Tõnisson, Karl Kramer, Katharina Götze, Florian Bassermann, Judith Schlegl, Hans-Christian Ehrlich, Stephan Aiche, Axel Walch, Philipp A Greif, Sabine Schneider, Eduard Rudolf Felder, Juergen Ruland, Guillaume Médard, Irmela Jeremias, Karsten Spiekermann, and Bernhard Kuster. The target landscape of clinical kinase drugs. Science, 358(6367):eaan4368, dec 2017.

[20] Stephen P Davies, Helen Reddy, Matilde Caivano, and Philip Cohen. Specificity and mechanism of action of some commonly used protein kinase inhibitors. Biochem. J, 351:95–105, 2000.

[21] Jenny Bain, Hilary Mclauchlan, Matthew Elliott, and Philip Cohen. The specificities of protein kinase inhibitors : an update. Biochem. J, 371:199–204, 2003.

[22] Jenny Bain, Lorna Plater, Matt Elliott, Natalia Shpiro, C. James Hastie, Hilary Mclauchlan, Iva Klevernic, Simon C. Arthur, Dario R. Alessi, and Philip Cohen. The selectivity of protein kinase inhibitors: a further update. Biochemical Journal, 408(3), 2007.

[23] Silvia Domcke, Rileen Sinha, Douglas A Levine, Chris Sander, and Nikolaus Schultz. Evaluating cell lines as tumour models by comparison of genomic profiles. Nature communications, 4, 2013.

[24] The Cancer Genome Atlas Network et al. Genomic classification of cutaneous melanoma. Cell, 161(7):1681–1696, 2015.

[25] The cbioportal for cancer genomics. http://www.cbioportal.org/patient?sampleId=A2058_SKIN&studyId=cellline_ccle_broad. Accessed: 2019-01-06.

[26] Eliezer M Van Allen, Nikhil Wagle, Antje Sucker, Daniel J Treacy, Cory M Johannessen, Eva M Goetz, Chelsea S Place, Amaro Taylor-Weiner, Steven Whittaker, Gregory V Kryukov, et al. The genetic landscape of clinical resistance to raf inhibition in metastatic melanoma. Cancer discovery, 4(1):94–109, 2014.

[27] F Xing, Y Persaud, CA Pratilas, BS Taylor, M Janakiraman, QB She, H Gallardo, C Liu, T Merghoub, B Hefter, et al. Concurrent loss of the pten and rb1 tumor suppressors attenuates raf dependence in melanomas harboring v600ebraf. Oncogene, 31(4):446–457, 2012.

[28] Lise Boussemart, Héléne Malka-Mahieu, Isabelle Girault, Delphine Allard, Oskar Hemmingsson, Gorana Tomasic, Marina Thomas, Christine Basmadjian, Nigel Ribeiro, Frédéric Thuaud, et al. eif4f is a nexus of resistance to anti-braf and anti-mek cancer therapies. Nature, 513(7516):105–109, 2014.

[29] Diederik P Kingma and Jimmy Ba. Adam: A method for stochastic optimization. arXiv preprint arXiv:1412.6980, 2014.

[30] R.D. Cook and S. Weisberg. Residuals and Influence in Regression. Monographs on statistics and applied probability. Chapman & Hall, 1982.

[31] Wenbin Liu, Zhenlin Ju, Yiling Lu, Gordon B Mills, and Rehan Akbani. A comprehensive comparison of normalization methods for loading control and variance stabilization of reverse-phase protein array data. Cancer informatics, 13:CIN–S13329, 2014.

[32] Stanley F Barnett, Deborah Defeo-Jones, Sheng Fu, Paula J Hancock, Kathleen M Haskell, Raymond E Jones, Jason A Kahana, Astrid M Kral, Karen Leander, Ling L Lee, John Malinowski, Elizabeth M McAvoy, Debbie D Nahas, Ronald G Robinson, and Hans E Huber. Identification and characterization of pleckstrin-homology-domain-dependent and isoenzyme-specific Akt inhibitors. The Biochemical journal, 385(Pt 2):399–408, 1 2005.

[33] Jochen Schust, Bianca Sperl, Angela Hollis, Thomas U. Mayer, and Thorsten Berg. Stattic: A Small-Molecule Inhibitor of STAT3 Activation and Dimerization. Chemistry & Biology, 13(11):1235–1242, 11 2006.

[34] Kenta Hara, Yoshiko Maruki, Xiaomeng Long, Ken-ichi Yoshino, Noriko Oshiro, Sujuti Hidayat, Chiharu Tokunaga, Joseph Avruch, and Kazuyoshi Yonezawa. Raptor, a Binding Partner of Target of Rapamycin (TOR), Mediates TOR Action. Cell, 110(2):177–189, 7 2002.

[35] Do-Hyung Kim, Dos D. Sarbassov, Siraj M. Ali, Jessie E. King, Robert R. Latek, Hediye Erdjument-Bromage, Paul Tempst, and David M. Sabatini. mTOR Interacts with Raptor to Form a Nutrient-Sensitive Complex that Signals to the Cell Growth Machinery. Cell, 110(2):163–175, 7 2002.

[36] J. Tsai, J. T. Lee, W. Wang, J. Zhang, H. Cho, S. Mamo, R. Bremer, S. Gillette, J. Kong, N. K. Haass, Sproesser, L. Li, K. S. M. Smalley, D. Fong, Y.-L. Zhu, A. Marimuthu, H. Nguyen, B. Lam, J. Liu, I. Cheung, J. Rice, Y. Suzuki, C. Luu, C. Settachatgul, R. Shellooe, J. Cantwell, S.-H. Kim, J. Schlessinger, K. Y. J. Zhang, B. L. West, B. Powell, G. Habets, C. Zhang, P. N. Ibrahim, P. Hirth, D. R. Artis, M. Herlyn, and G. Bollag. Discovery of a selective inhibitor of oncogenic B-Raf kinase with potent antimelanoma activity. Proceedings of the National Academy of Sciences, 105(8):3041–3046, 2 2008.

[37] Tinghu Zhang, Francisco Inesta-Vaquera, Mario Niepel, Jianming Zhang, Scott B Ficarro, Thomas Machleidt, Ting Xie, Jarrod A Marto, NamDoo Kim, Taebo Sim, John D Laughlin, Hajeung Park, Philip V LoGrasso, Matt Patricelli, Tyzoon K Nomanbhoy, Peter K Sorger, Dario R Alessi, and Nathanael S Gray. Discovery of potent and selective covalent inhibitors of JNK. Chemistry & biology, 19(1):140–54, 1 2012.

